# Visualizing ATP-dependent substrate-processing dynamics of the human 26S proteasome at near-atomic resolution

**DOI:** 10.1101/419051

**Authors:** Yuanchen Dong, Shuwen Zhang, Zhaolong Wu, Xuemei Li, Wei Li Wang, Yanan Zhu, Svetla Stoilova-McPhie, Ying Lu, Daniel Finley, Youdong Mao

## Abstract

The proteasome is an ATP-dependent 2.5-megadalton machine responsible for ubiquitylated protein degradation in all eukaryotic cells. Here we present cryo-EM structures of the substrate-engaged human 26S proteasome in seven conformational states at 2.8-3.6 Å resolution, captured during polyubiquitylated protein degradation. These structures visualize a continuum of dynamic substrate-proteasome interactions from ubiquitin recognition to processive substrate translocation, during which ATP hydrolysis sequentially navigate through all six ATPase subunits. Three principle modes of coordinated ATP hydrolysis are observed, featuring hydrolytic events in two oppositely positioned ATPases, in two consecutive ATPases, and in one ATPase at a time. They regulate deubiquitylation, translocation initiation and processive unfolding of substrates, respectively. A collective power stroke in the ATPase motor is generated by synchronized ATP binding and ADP release in the substrate-engaging and disengaging ATPases, respectively. It is amplified largely in the substrate-disengaging ATPase, and propagated unidirectionally by coordinated ATP hydrolysis in the third consecutive ATPase.

---

The ubiquitin-proteasome pathway (UPP) plays a central role in selective protein degradation in all eukaryotic cells. It is involved in the regulation of myriad cellular processes, such as cell cycle, apoptosis, immune response and inflammation, neural and muscular degeneration, the response to proteotoxic stress, and so fourth^1-5^. The UPP operates through two discrete, successive processes: (1) tagging of the target substrates through numerous pathways of ubiquitylation^6^, and (2) recognition and degradation of the tagged substrates by the 2.5-MDa 26S proteasome holoenzyme^7^. The proteasome holoenzyme is assembled from a barrel-shaped, proteolytically active core particle (CP), also called the 20S proteasome, and two 19S regulatory particles (RP)^8-10^ capping at both ends of the CP cylinder. The RP controls the substrate access into the CP^11-16^ and is formed from two subcomplexes known as the lid and the base. Recognition of a ubiquitylated substrate is mediated principally by ubiquitin receptors Rpn1, Rpn10 and Rpn13 of the base^4,17^. Once captured by the RP, the globular domains of a substrate, whose conjugated ubiquitin chains are first removed by a metalloprotease lid subunit, Rpn11, and then mechanically unfolded by a ring-like heterohexameric motor module in the base^18^. The motor module consists of six distinct adenosine triphosphatase (ATPase) subunits, Rpt1-6, which belong to the ATPases-associated-with-diverse-cellular-activities (AAA) family. The AAA-ATPase motor regulates the engagement, deubiquitylation and degradation of substrates in an ATP-dependent manner^19-21^. The exact mechanism of such regulations is the most enigmatic open problem in our understanding of the 26S proteasome function^22^.

Previous structural studies with cryo-electron microscopy (cryo-EM) have revealed the general architecture of the substrate-free holoenzyme in six distinct states^15,16,23-29^, which provides a foundation to appreciate how the 26S proteasome dynamically regulates its functions. However, it remains unclear how the conformations of the substrate-free proteasome holoenzyme are related to their functional states in the presence of a substrate. The mechanism by which the substrate is engaged, deubiquitylated, unfolded and translocated by the 26S proteasome remains elusive due to lack of high-resolution structures of the substrate-bound proteasomes. In this work, we have captured the substrate-engaged human 26S proteasome during ATP-dependent protein degradation and determined cryo-EM structures of the human 26S proteasome in complex with the model substrate Sic1^PY^ in seven conformational states at nominal resolutions of 2.8-3.6 Å (Fig. 1, Extended Data Figs. 1-3, Extended Data Table 1)^30^.

**Figure 1.**
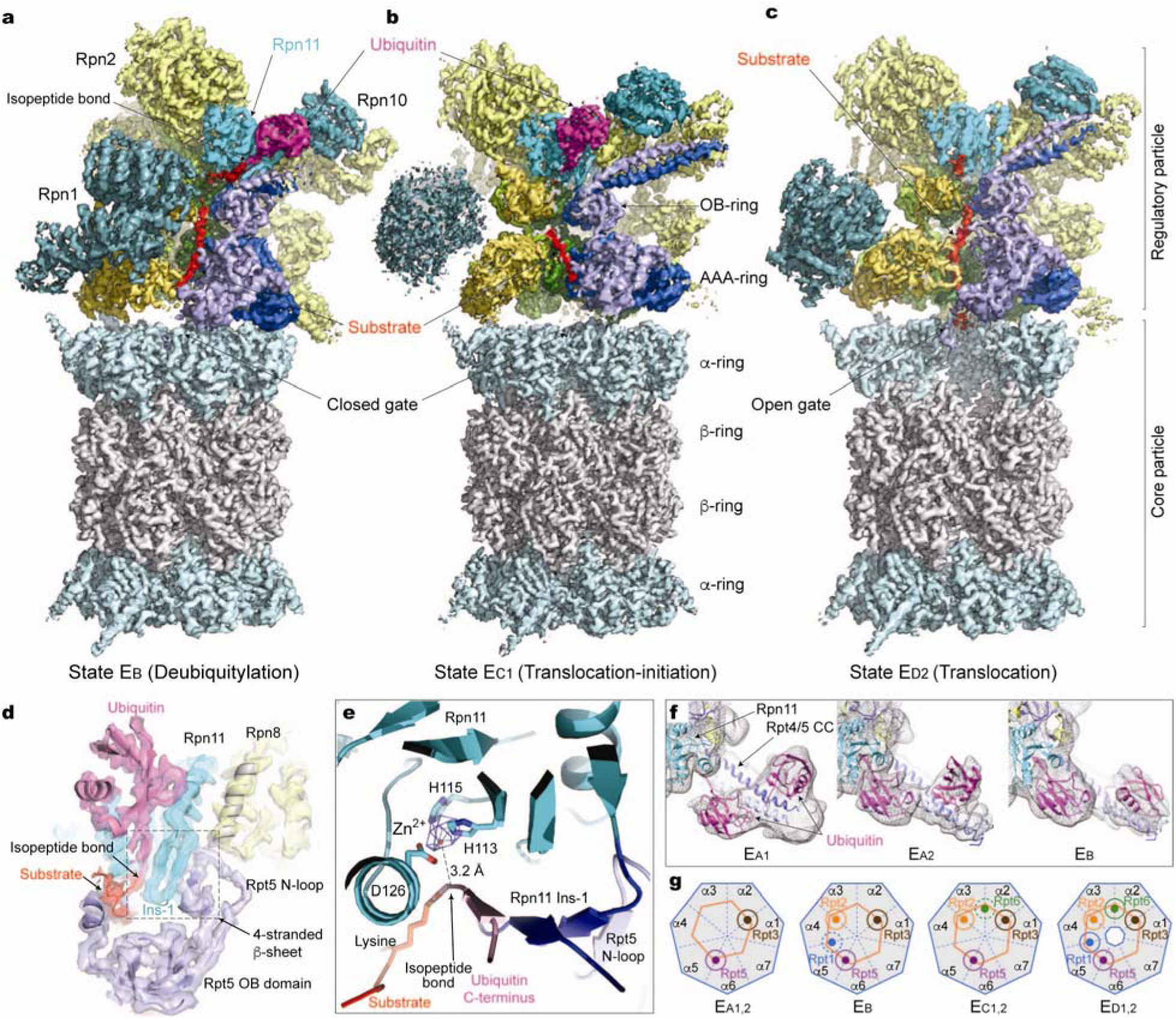
Cryo-EM structures of the substrate-bound human 26S proteasome in distinct states. (**a**) Cryo-EM density map of the substrate-bound human 26S proteasome in state E_B_ at 3.3 Å shows a ubiquitin-conjugated substrate (red) is engaged with the AAA-ATPase channel, with the ubiquitin bound to Rpn11 and the isopeptide bond lining near the zinc-binding site. (**b**) Cryo-EM density map of the substrate-bound human 26S proteasome in state E_C1_ at 3.5 Å shows that the ubiquitin binds on the Rpn11 but is no longer covalently linked to the substrate. (**c**) Cryo-EM density map of the substrate-bound human 26S proteasome in state E_D2_ at 3.2 Å shows that the ubiquitin is gone, and the substrate is threading into the open CP gate. The Rpt1 density is omitted in (a-c) in order to show the substrate density inside the AAA-ATPase. Two α-subunits are omitted in order to show the substrate density inside the CP gate in (c). (**d**) A close-up view of the quaternary interface around the isopeptide bond between ubiquitin and the substrate lysine in state E_B_. The scissile isopeptide bond density is visible at the present resolution and is labeled. The cryo-EM density in state E_B_ is rendered as a transparent surface, superimposed with the cartoon representation of the atomic model. The C-terminal strand of ubiquitin, the Insertion-1 loop and the C-terminal strand of Rpt5 N-loop form a quaternary four-stranded β-sheet, with the Rpt5 N-loop filling up the cavity between Rpn11 Insertion-1 and Rpn8. (**e**) A close-up view of the zinc ion (hotpink sphere) immediately approached by the isopeptide bond between the ubiquitin C-terminus and the substrate lysine. The zinc ion density is shown as a blue mesh at 10σ level. The quaternary four-stranded β-sheet supports the interaction between the isopeptide bond and the zinc ion. The view is rotated 90° relative to (d). Three side chains of Rpn11 coordinating with the zinc ion are shown and labeled. (**f**) Comparison of two ubiquitin moieties between Rpn11 and Rpt4/5 CC among states E_A1_, E_A2_ and E_B_. The cryo-EM densities of the three states rendered as grey mesh representations are low-pass filtered to 8 Å in order to show the second ubiquitin density at a low resolution. The atomic model of ubiquitin is shown as a magenta cartoon representation. Both ubiquitins reside on the N-terminal portion of the Rpt4/5 CC and are not in direct contact with Rpn11 in state E_A1_. One of them is in contact with both Rpn11 and Rpt4/5 CC in state E_A2_, but only Rpn11 in state E_B_. (**g**) A schematic diagram summarizes the correlation of the Rpt C-terminal insertion to the α-pockets of the CP and the state of CP gate in the seven states. The CP is represented as a heptagon, the AAA-ATPase as a hexagon, and the Rpt C-tail insertion as a colored sphere in a circle.

These cryo-EM structures together visualize a continuum of dynamic substrate-proteasome interactions, from initial ubiquitin recognition^31,32^, to ATP-dependent deubiquitylation^33^, and to processive substrate translocation. Importantly, we observed eight events of ATP hydrolysis sequentially navigating throughout all six ATPase subunits. Our analysis reveals three principle modes of coordinated ATP hydrolysis regulating different stages of substrate processing, and provides comprehensive insights into the complete cycle of substrate degradation by the human proteasome holoenzyme.

## Overview of the conformational states

States E_A1,2_ are overall similar to the substrate-free S_A_ conformation of human proteasome holoenzyme in their CP and ATPase components (Extended Data Fig. 3)^15,23,24,26^. Several remarkable features distinguish states E_A1,2_ from S_A_. Foremost, a ubiquitin density is observed on the T1 site of Rpn1, and two ubiquitin densities at the N-terminal segment of Rpt4-Rpt5 N-terminal coiled coil (CC) next to Rpn10 (Fig. 1f and Extended Data Fig. 4a)^17^. States E_A1_ and E_A2_ capture the snapshots before and after one ubiquitin near Rpn10 is engaged with Rpn11, respectively. States E_A1,2_ also present 13 potential substrate densities in the chamber of the closed CP, including one bound to the proteolytic site Thr1 of the β2 subunit (Extended Data Fig. 5a and Extended Data Table 2). However, no substrate is found inside the AAA-ATPase channel (Fig. 2a). Therefore, state E_A_ presumably represents the conformation of initial ubiquitin recognition, albeit after the holoenzyme has been recycled from a previous degradation event.

**Figure 2.**
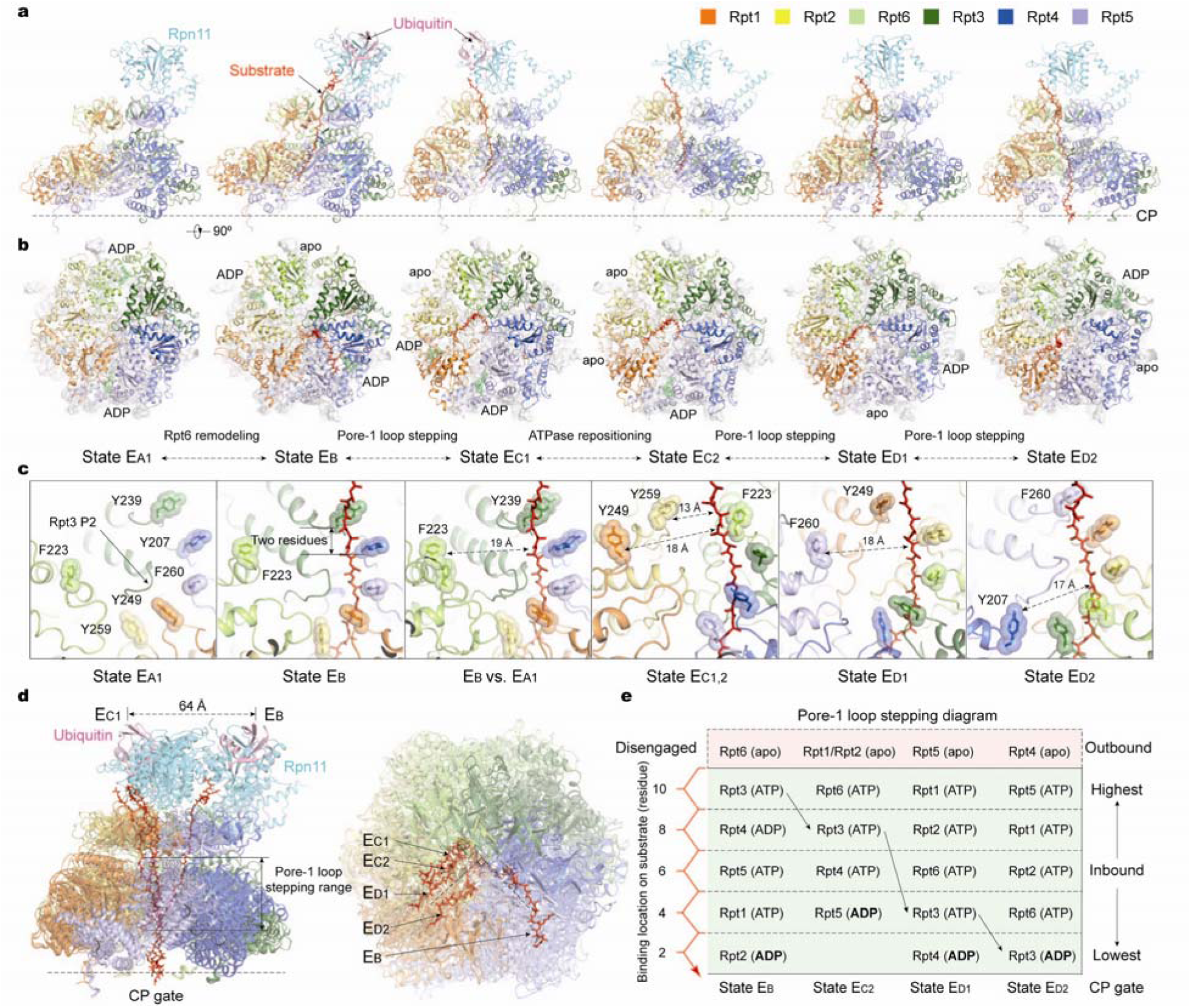
Dynamic substrate-proteasome interactions. (**a**) Side views of the AAA-ATPase-Rpn11 subcomplex interacting with the substrate in five states (E_B_, E_C1,2_ and E_D1,2_) in comparison with state E_A1_. The substrate is modelled as a polypeptide backbone structure and is represented as red sticks. Ubiquitin, Rpn11 and AAA-ATPase are rendered as transparent cartoons to show the substrate translocating inside the central channel. The relative location of the CP is marked by the horizontal dashed line. (**b**) Top views of the AAA-ATPases of the same states shown in (a). Both ADP-bound and apo-like Rpt subunits are labeled, where the remaining Rpt subunits are all bound with ATP. Nucleotides are shown as stick representations. The stick representation of ADP is colored green and is superimposed with green transparent sphere representation. The CP is rendered as a grey surface representation. The seven structures in (a) and (b) are aligned altogether only using their CP components to show the relative motion of their AAA-ATPases against the common CP. (**c**) The varying architecture of pore-1 loop staircase in direct contact with the substrate in five states (E_B_, E_C1,2_ and E_D1,2_) as compared to that in state E_A1_. The aromatic residues in the pore-1 loops are labeled and shown as stick representations superposed with transparent sphere representations for highlighting. In all case, one or two pore-1 loops are disengaged with the substrate-pore loop staircase, and their distances to the substrate are marked. The third panel shows the superposition of state E_B_ with state E_A_ to illustrate the similarity of the pore-1 loop architecture before and after the substrate engagement. The distance between two nearest pore-1 loop contacts invariantly spans across two amino acids. The AAA-ATPase structures in states E_C1_ and E_C2_ remains nearly identical except for different nucleotide states and their relative position above the CP. Thus, only one structure is shown in the fourth panel to represent both. (**d**) The side view (left) and top view (right) of the seven structures shown in (a) and (b) are superimposed together based on the structure alignment against the CP, showing how the substrate-translocation pathway is significantly changed from state E_B_ to states E_C/D_. (**e**) A diagram summarizing the variation of the pore-1 loops interacting with the substrate observed in states E_B/C/D_. The vertical axis shows the relative location of pore-1 loops interacting the substrate, with the CP positioned at the bottom. The numbers label the relative distance from the lowest substrate-pore loop contact, using the number of residues as a metric. The horizontal axis shows the four states with different substrate-pore loop architectures. State E_C1_ is omitted here due to its AAA-ATPase structure being identical to state E_C2_. In all four states, the ADP is bound to the lowest Rpt subunits and the apo-like subunits are disengaged from the substrate. The color code for subunit structures in all panels are shown in the upper right corner.

State E_B_ reveals a novel deubiquitylation-compatible complex that does not resemble any known substrate-free proteasome conformation (Fig. 1a)^26^. Most notably, the isopeptide bond between substrate lysine and ubiquitin is visible in the vicinity to the zinc-binding site of Rpn11, through extensive quaternary rearrangements (Fig. 1d). To facilitate such a quaternary rearrangement, the lid is concomitantly rotated outward away from the axis of the oligonucleotide-/oligosaccharide-binding (OB) ring in the base relative to state E_A_, to a direction opposite to that seen in the E_A_-to-E_D_ transition (Fig. 2a, Extended Data Fig. 4d,e)^34,35^. This gives rise to a more open entrance to the AAA-ATPase central channel. Although four C-terminal tails from Rpt1, Rpt2, Rpt3 and Rpt5 are inserted into the inter-subunit surface pockets of α-subunits, the CP gate is closed (Fig. 1g, Extended Data Fig. 4g).

E_C1_ and E_C2_ presents two successive states that are compatible with the initiation of substrate translocation (Figs. 1b,2a,b, Extended Data Fig. 3). Ubiquitin remains on Rpn11 in state E_C1_; but interestingly it is released in state E_C2_. However, the two states exhibit similar features in the Rpt ring and CP gate. In addition to three Rpt C-terminal HbYX motifs inserted in the α-pockets^36^, weak densities are observed for the Rpt6 C-terminal tail in the α2-α3 pocket in both states (Fig. 1g, Extended Data Fig. 4g). The CP gate is still closed. The ATPase ring exhibits a rigid-body motion above the CP, with a small rotation in the lid. The substrate in the AAA-ATPase channel is hypothetically advanced forward relative to E_B_. The overall lid-base relationship resembles state E_D_.

States E_D1_ and E_D2_ capture two consecutive conformations in which distinct substrate-pore loop contacts are compatible with processive substrate translocation (Figs. 1c,2a,b, Extended Data Fig. 3). The substrate is presumably advanced forward relative to E_C1,2_. No ubiquitin densities are observed in these states. The CP gate is opened by the insertion of the C-terminal tails of five Rpt subunits, except Rpt4, into the α-pockets^26^. The general lid-base relationship and the open state of CP gate resembles the S_D_-like conformations of the substrate-free human proteasome^15,26^.

## Quaternary rearrangement for substrate deubiquitylation

A prominent feature in state E_B_ is the formation of a quaternary subcomplex involving substrate-ubiquitin-bound Rpn11, Rpn8 and Rpt5 N-loop (Fig. 1d-f). This quaternary subcomplex already starts to form in state E_A2_, where the AAA-ATPase is not yet engaged with the substrate (Extended Data Fig. 4a-c). The Rpn11-bound ubiquitin also contacts the Rpt4-Rpt5 N-terminal CC in E_A2_ and resides midway between its positions in states E_A1_ and E_B_ (Fig. 1f). The lid is rotated toward the same direction as that in state E_B_ (Extended Data Fig. 4d,e). The Rpt4-Rpt5 CC is, however, shifted up, allowing the gap between Rpn11 and Rpt4-Rpt5 CC to shrink. In state E_A1_, this ubiquitin resides on the Rpt4-Rpt5 CC in the vicinity of Rpn11 but does not contact Rpn11 (Fig. 1f). This supports our speculation that E_A2_ captures an intermediate state where a ubiquitin is being transferred from Rpt4-Rpt5 CC to Rpn11.

The Rpn11-ubiquitin interface is centered on a hydrophobic pocket around Trp111 and Phe133 of Rpn11, with the positioning of ubiquitin being identical to that observed in the crystal structure of the ubiquitin-bound Rpn11-Rpn8 complex^33^. The Insertion-1 loop of Rpn11 assumes a β-hairpin conformation, and pairs from both sides with the C-terminal strand of ubiquitin and a segment of the Rpt5 N-loop emanating from the OB ring, forming a four-stranded β-sheet (Fig. 1d). This quaternary β-sheet directs the ubiquitin C-terminal toward the proteolytically active zinc-binding site in Rpn11 and places the isopeptide bond right next to the zinc ion, which has a strong density in our cryo-EM maps (Fig. 1e). Compared to the crystal structure of ubiquitin-bound Rpn11, the Insertion-1 β-hairpin is tilted outward in state E_B_ and is better aligned with the zinc-binding site in the proteasome^33^. This finding suggests that the conserved Rpt5 N-loop stabilizes the Rpn11-ubiquitin quaternary contact and optimizes the orientation of the isopeptide bond for efficient deubiquitylation. By contrast, the Rpt5 N-loop is disordered in most other states (E_A1_, E_C1,2_ and E_D1,2_).

The Insertion-1 loop of Rpn11 alternates among three distinct configurations through all seven states (Extended Data Fig. 4c). The Insertion-1 loop remains as a β-hairpin throughout states E_A2_, E_B_ and E_C1_, whenever ubiquitin is bound. By contrast, it is a large open loop in state E_A1_, and refolds into a smaller, tighter loop in states E_C2_, E_D1,2_, where ubiquitin is released. Taken together, the quaternary organization surrounding the zinc-binding site and the extended interactions between ubiquitin chains and the quaternary structure of Rpn10-Rpn11-Rpt4-Rpt5 appear to explain why the Rpn11 exhibits much higher deubiquitylation activity in the proteasome context than in its non-proteasome forms, such as in the Rpn8-Rpn11 dimeric form^37,38^, the free lid subcomplex^39^ and the free RP assembly^40^.

## Substrate interactions with the holoenzyme

In the progression from states E_B_ to E_D1,2_, the substrate contact with Rpn11 is invariantly centered around a hydrophobic groove at Phe118 and Trp121 of Rpn11. In state E_B_, this binding site faces the Rpt3-Rpt4 OB interface and the substrate extends straight from this site to Val125 of Rpn11, beneath which the isopeptide bond linking the ubiquitin with the substrate is held. In states E_D1,2_, with the CP gate open and the ubiquitin released, the substrate is free to thread through the AAA-ATPase channel. Inside the constriction of the OB ring, the substrate density is well centered in state E_B_, without direct contact with the interior surface of the ring. This approximate symmetry is broken in states E_D1,2_, where the substrate closely approaches Phe118 of Rpt1 in the interior of the OB ring.

Within the central channel of the AAA ring, the substrate is gripped into a right-handed spiral staircase architecture in direct contact with the aromatic residues of pore-1 loops (Fig. 2c, Extended Data Fig. 5b-d). The aromatic residues, either tyrosine or phenylalanine, intercalate with the zigzagging main chain through hydrophobic interactions. In addition, the main chains of the pore-1 loops potentially form hydrogen bonds with the main chain of substrate. The pore-1 loops are evenly distributed along the substrate, with two adjacent pore-1 loops spanning two consecutive amino acid residues in the substrate. This substrate-pore loop staircase architecture is nearly invariant from states E_B_ to E_D2_ (Fig. 2a-d). The pore-1 loop staircase in E_B_ is highly similar to that in E_A_, suggesting that E_B_ is an initially substrate-engaged state before any translocation occurs, consistent with the observation of the isopeptide bond linking substrate with ubiquitin on Rpn11 in E_B_. The pore-1 loops of Rpt3, Rpt6, Rpt1 and Rpt5 reside at the highest position in contact with the substrate in states E_B_, E_C_, E_D1_ and E_D2_, respectively. Notably, the pore-1 loop of Rpt3 moves from the highest position to the lowest position in the substrate-bound pore-loop staircase from E_B_ to E_D2_. Meanwhile, the pore-2 loops support the opposite side of the substrate through their charged acidic residues, forming another, somewhat shorter staircase beneath the pore-1 loop staircase. In states E_B_, E_D1_ and E_D2_, the pore loops from Rpt6, Rpt5 and Rpt4 are disengaged from the substrate (Fig. 2c). In contrast, the pore loops from both Rpt1 and Rpt2 are disengaged from the substrate in E_C1,2_.

## A continuum of nucleotide state transitions

The current resolution allows us to unambiguously distinguish ADP from ATP in the nucleotide-binding pockets of the ATPases in most cases (Fig. 2b,e, Methods and Extended Data Figs. 3c,6). Except for E_A_, at least one ATPase subunit in each conformational state exhibits a very weak or partial density for a potential nucleotide in its nucleotide-binding pocket, which precludes *de novo* atomic modeling of nucleotide coordinates into the density of that subunit. We refer to the nucleotide state of these ATPases as an apo-like state. It is not possible to differentiate between ATP and ATP-γS at the present resolution, given a mixture of both ATP and ATP-γS in the protein buffer (see Methods). For simplicity, we will refer to nonhydrolyzed nucleotide as ATP.

Notably, the ADP-bound states navigate counterclockwise sequentially from Rpt6 to Rpt3 throughout all six ATPase subunits, indicating a full cycle of coordinated ATP hydrolysis throughout the AAA-ATPase ring from state E_A_ to E_D2_ (Fig. 2b). However, eight events of coordinated ATP hydrolysis are observed in total, because two ADPs were observed in two oppositely positioned ATPases in states E_A_ and E_B_. In state E_A_, Rpt5 and Rpt6 are both loaded with ADP, whereas the other four ATPases are all loaded with ATP. Magnesium ion density is evident next to ATP but not ADP in the E_A_ density map (Extended Data Fig. 3c). In state E_B_, Rpt6 assumes the apo-like state, whereas Rpt2 and Rpt4, the nearest counterclockwise neighbors to the two ADP-bound subunits in E_A_, are bound with ADP (Fig. 2b, Extended Data Fig. 6). The AAA domain of Rpt6 is displaced away from the rest of the AAA ring, forming prominent gaps at the Rpt2-Rpt6 and Rpt6-Rpt3 interfaces (Fig. 3a).

**Figure 3.**
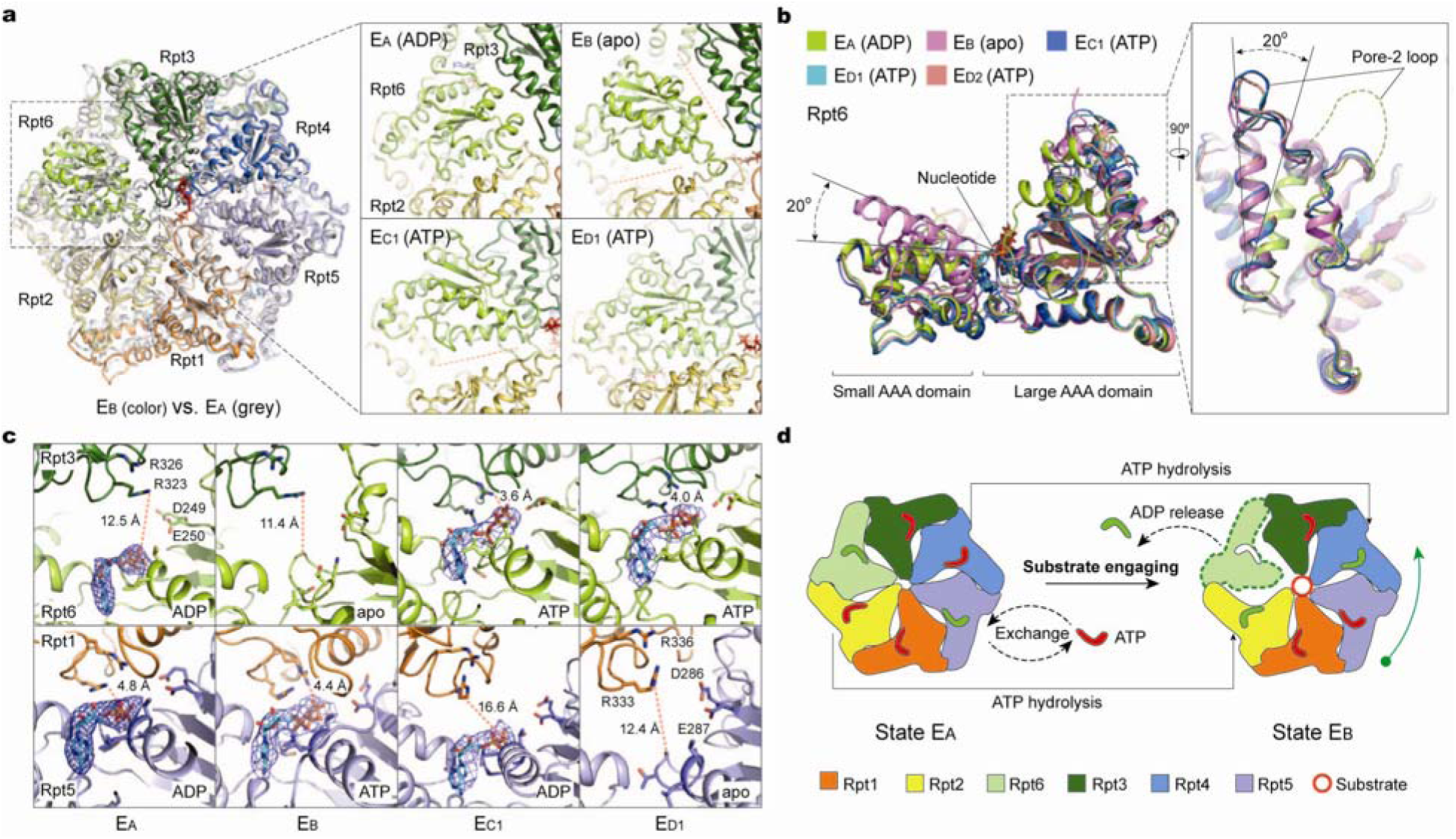
Structural basis for nucleotide-driven substrate engagement in the AAA-ATPase channel. (**a**) Superposition of the AAA-ring structures of states E_A_ (grey) and E_B_ (color) shows that only Rpt6 exhibits a large-scale conformational rearrangement during E_A_-to-E_B_ transition, whereas other five Rpt subunits only make subtle local adjustments. The insets show side-by-side comparison of the Rpt6 conformations in four most distant states. Two large gaps are created at Rpt2-Rpt6 and Rpt6-Rpt3 interfaces in state E_B_. The Rpt6-Rpt3 gap is closed in state E_C1_. The Rpt2-Rpt6 gap remains open in state E_C1_ and is closed in state E_D1_. (**b**) Superposition of the Rpt6 AAA domain structures from five distinct states aligned against the large AAA subdomain shows that Rpt6 assumes three major conformations. Transition from state E_A_ to E_B_ involves both refolding of the pore-2 loop, shown in the right insert, and a 20° rigid-body rotation between the large and small AAA subdomains. Transition from state E_B_ to E_C,D_ involves only the rigid-body rotation between the large and small AAA subdomains. The ATP-bound Rpt6 structures are nearly all identical in different states. (**c**) Comparison of the nucleotide-binding pockets of Rpt6 and Rpt5 in four states illustrates a common pattern of the openness of the nucleotide-binding sites. The cryo-EM densities of the nucleotides are shown as blue meshes. When the Rpt is in the middle of pore-loop staircase but not at the lowest position, the nucleotide-binding pockets are closed no matter it is ATP or ADP bound. By contrast, the nucleotide-binding pocket is widely open no matter if it is ADP bound or free of nucleotide when the Rpt is either in the lowest position of the pore-loop staircase or disengaged from the staircase. (**d**) A cartoon formula summarizes the ATP hydrolysis reactions and nucleotide exchange events required to implement the state transition from E_A_ to E_B_, which results in the substrate engagement and deubiquitylation. Panels (a), (c) and (d) share the same subunit color code.

The nucleotide states or the geometry of the nucleotide-binding pockets of the ATPases are strongly correlated with their substrate-pore loop interactions (Fig. 2e). The apo-like state is always observed in the ATPases whose pore loops are disengaged from the substrate. Thus, all apo-like subunits form prominent gaps at the interfaces with their nearest neighbors on both sides, resembling the case of Rpt6 in state E_B_ (Figs. 2b,3a,c). The nucleotide-binding pockets of the apo-like subunits consistently exhibit prominent openness, with the arginine fingers from the clockwise adjacent subunit displaced more than 10 Å away from the Walker A motif (Fig. 3c). The ATPase whose pore-1 loop resides at the lowest position in direct contact with the substrate is always bound with ADP. Thus, the counterclockwise nearest neighbor of the apo-like subunit is found to be ADP-bound in all cases (Fig. 2b,3c). Similarly, the nucleotide-binding pocket is always tightly closed, with the arginine fingers residing within 3-5 Å away from either γ-phosphate or β-phosphate, whenever the ATPases engaged with the substrate have their pore-1 loops located in the middle or top registry in the substrate-pore loop staircase. Except states E_A_ and E_B_, ATP is invariantly found to bind the substrate-engaged ATPases at the middle or top positions in the pore-loop staircase.

## ATP-dependent regulation of substrate engagement

Structural comparison between states E_A_ and E_B_ sheds lights on the question of how ATP hydrolysis in AAA-ATPases regulates the substrate engagement for deubiquitylation (Fig. 3). In state E_A_, the AAA channel is too narrow to engage the substrate, suggesting that it is in a closed state. To open the channel for the substrate engagement, the AAA-ATPase ring must rearrange its quaternary organization. Indeed, the AAA domain of Rpt6 undergoes dramatic structural changes from E_A_ to E_B_. An out-of-plane rotation of about 40° is observed in the large AAA subdomain of Rpt6, whereas the AAA domains of other ATPases mostly move as a rigid body with subtle changes restricted to the pore loops (Fig. 3a, b). In state E_A_, the pore-2 loop of Rpt6 is largely disordered; by contrast, the pore-2 loop refolds into a fairly ordered structure in E_B_, spanning residues 251-256 (Fig. 3b). Consistent with this observation, the ADP bound to Rpt6 in state E_A_ is released in E_B_. Thus, ATP hydrolysis and ADP release in Rpt6 are programed to trigger an iris-like movement in the AAA ring that opens the central channel (Fig. 3c, d). The coordinated ATP hydrolysis in Rpt5 directly opposite from Rpt6 in state E_A_, and in Rpt4 opposite from Rpt2 in state E_B_, is expected to increase the conformational flexibility of the AAA ring required to open the AAA channel for accommodating the substrate engagement (Fig. 3d). The position of Rpt6 in the AAA ring, which is disengaged from the pore-loop staircase in both states E_A_ and E_B_, endows it with certain energetic advantage in triggering the AAA channel opening with little perturbation to the pore loop staircase that is poised for accepting an unfolded terminus of the substrate without changing the overall architecture.

## Initiation of substrate translocation

From state E_A_ to E_D2_, the Rpt1-Rpt2 dimer and Rpt5 undergo a complete cycle of ATP hydrolysis and nucleotide exchange that regulates substrate translocation (Figs. 4a,5a). To allow a forward sliding process, at least one Rpt subunit in direct contact with the substrate at the lowest position must disengage from the substrate-pore loop staircase. Indeed, during the E_B_-to-E_C1_ transition, the pore loops of Rpt1-Rpt2 are disengaged from the substrate, partly facilitated by coordinated ATP hydrolysis in Rpt1 and Rpt5 as well as ADP release in Rpt2 (Fig. 4a, b). This is because ATP hydrolysis in Rpt5 would abolish the interaction of γ-phosphate with the trans-acting arginine finger from Rpt1, thus destabilizing the Rpt1-Rpt5 association. A large out-of-plane rotation of about 40° in the Rpt1-Rpt2 dimer moves the corresponding pore loops to almost the highest altitude but 13-18 Å away from the substrate (Figs. 2c,4a,b). Concomitantly, ATP binding to Rpt6 promotes the engagement of Rpt6 pore-1 loop with the substrate, resulting in one-step forward translocation of the substrate by a distance of two residues. The ADP release in Rpt1 during the E_C1_-to-E_C2_ transition does not trigger obvious conformational changes. However, during the E_C2_-to-E_D1_ transition, ATP binding to both Rpt1 and Rpt2, which drives their substrate re-engagement at the top of the substrate-pore loop staircase, gives rise to a small out-of-plane rotation of 5° in Rpt1-Rpt2, resulting in two-step forward translocation of the substrate (Fig. 4a,d,e). Similarly, the ATP-regulated substrate re-engagement of Rpt5 during the E_D1_-to-E_D2_ transition gives rise to substrate translocation by another step forward (Fig. 5).

**Figure 4.**
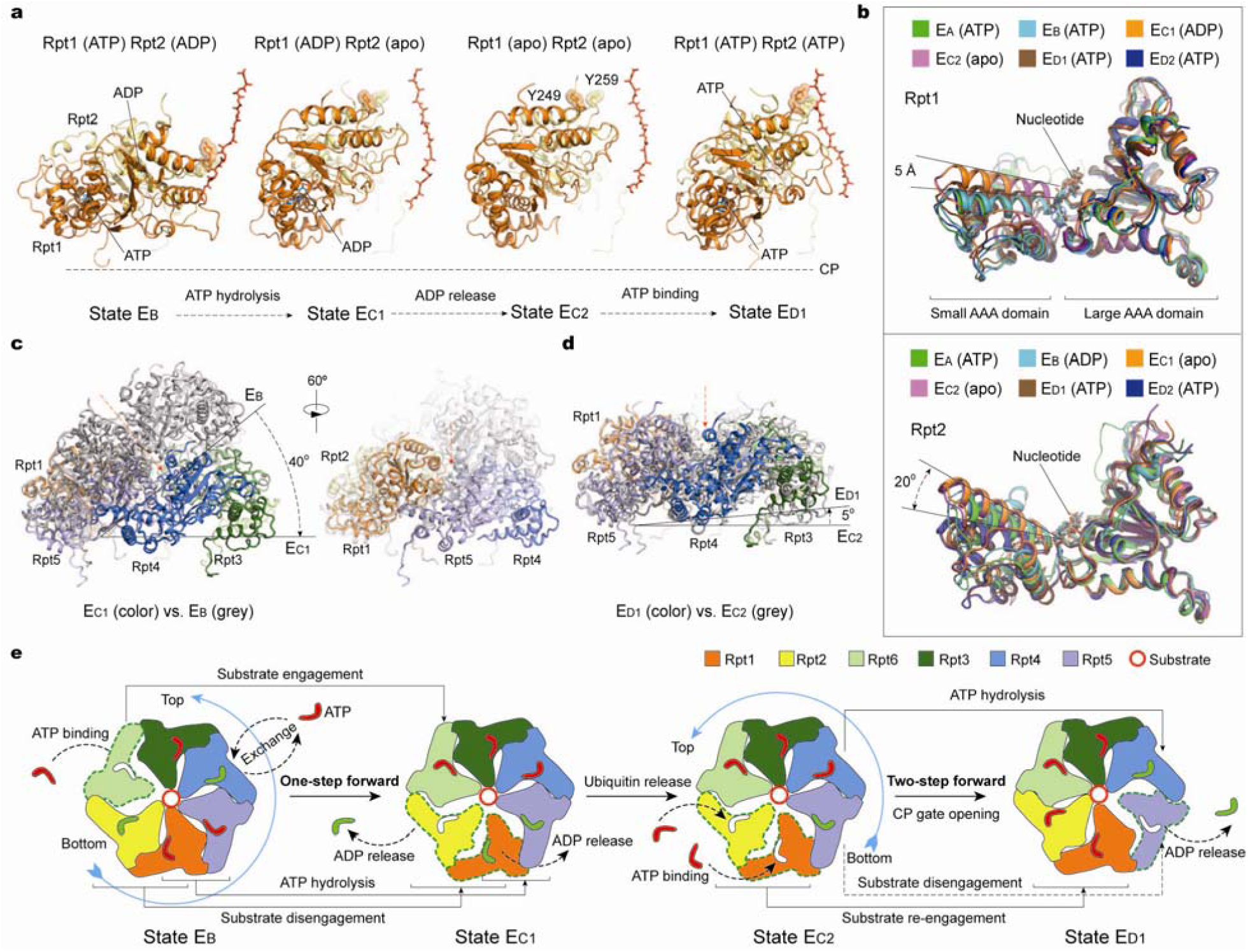
ATP hydrolysis drives initial three steps of substrate translocation in the AAA-ATPase channel. (**a**) Side-by-side comparison of the Rpt1-Rpt2 dimer conformations in four sequential states that cover a complete cycle of ATP hydrolysis and exchange in the Rpt1-Rpt2 dimer. The structures are aligned against the CP to show their conformational changes relative to the CP gate. The tyrosine residues in the pore-1 loops are shown as the stick representation and highlighted with the transparent sphere representation. Substrate is shown as the red stick representation. (**b**) Superposition of the Rpt1 (upper) or Rpt2 (lower) structures from six distinct states aligned against their large AAA subdomains shows that Rpt1 assumes two major conformations, and that Rpt2 assumes three major conformations. (**c**) Side views of the structure comparison of the AAA ring between E_C1_ (color) and E_B_ (grey), by using the large AAA subdomain of Rpt1 to align the two AAA-ring structures together, shows that a 40° out-of-plane rotation of the large AAA subdomain of Rpt1 relative to the AAA ring during the Rpt1-Rpt2 disengagement from the substrate. The right panel, rotated vertically against the left panel, shows that the out-of-plane rotation in Rpt1 is significantly amplified in its counterclockwise neighboring ATPases more than its clockwise neighbors. Red arrows mark the center of the AAA ring. (**d**) Structure comparison between E_D1_ (color) and E_C2_ (grey), by using the large AAA subdomain of Rpt1 to align the two AAA-ring structures together, shows that a small 5° out-of-plane rotation of the large AAA subdomain of Rpt1 relative to the AAA ring during the Rpt1-Rpt2 re-engagement with the substrate. (**e**) A cartoon formula summarizes the cascade of ATP hydrolysis reactions and nucleotide exchange events required to implement the state transitions from E_B_ to E_D1_, which results in substrate translocation by three steps in total. Panels (a) and (c-e) share the same subunit color code.

**Figure 5.**
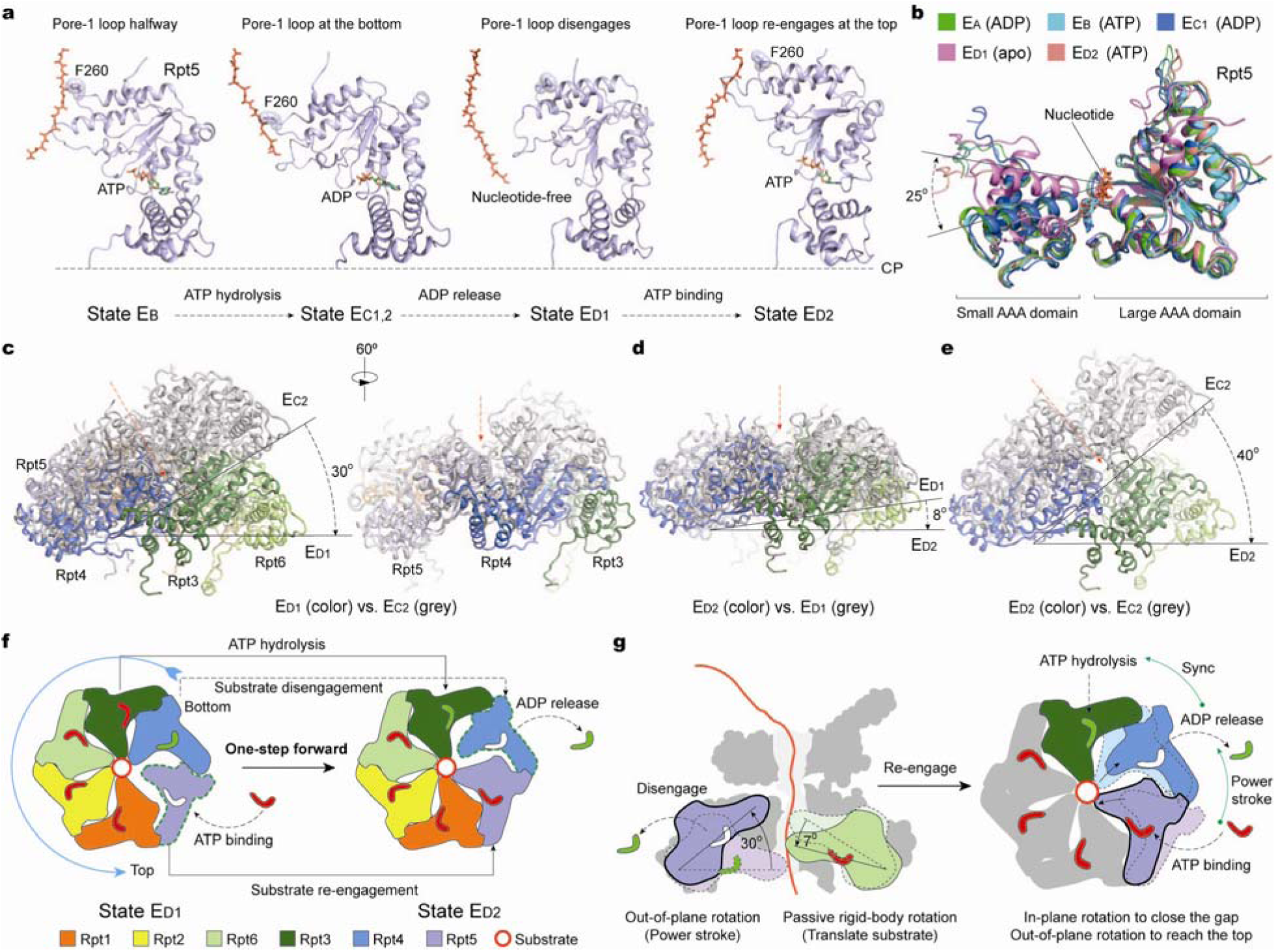
Mechanism for processive substrate translocation driven by a complete cycle of ATP hydrolysis. (**a**) Side-by-side comparison of the Rpt5 conformations in four sequential states that cover a complete cycle of ATP hydrolysis and exchange in Rpt5. The structures are aligned against the CP to show their conformational changes relative to the CP gate. The tyrosine residue in the pore-1 loop is shown as the stick representation and highlighted with the transparent sphere representation. Substrate is shown as the red stick representation. (**b**) Superposition of the Rpt5 structures from five distinct states aligned against the large AAA subdomain shows that Rpt5 assumes two major conformations. (**c**) Structure comparison between E_D1_ (color) and E_C2_ (grey), by using the large AAA subdomain of Rpt5 to align the two AAA-ring structures together, shows that a 30° out-of-plane rotation of the large AAA subdomain of Rpt5 relative to the AAA ring during Rpt5 disengagement from the substrate. The right panel, rotated vertically against the left panel, shows that the out-of-plane rotation in Rpt5 is significantly amplified in its counterclockwise neighboring ATPases more than its clockwise neighbors. Red arrows mark the center of the AAA ring. (**d**) Structure comparison between E_D2_ (color) and E_D1_ (grey), by using the large AAA subdomain of Rpt5 to align the two AAA-ring structures together, shows that a 8° out-of-plane rotation of the large AAA subdomain of Rpt5 relative to the AAA ring during Rpt5 re-engagement with the substrate. (**e**) Structure comparison between E_D2_ (color) and E_C2_ (grey), using the same alignment procedure, shows that a collective out-of-plane rotation of 40° in the large AAA subdomain of Rpt5 relative to the AAA ring during the cycle of Rpt5 disengagement from and reengagement with the substrate. (**f**) A cartoon formula summarizes the ATP hydrolysis reactions and nucleotide exchange events required to implement the state transitions from E_D1_ to E_D2_, which results in substrate translocation by one step. Panels (a) and (c-f) share the same subunit color code. (**g**) A cartoon diagram illustrates a model for the major power stoke generated during the disengagement of the Rpt subunit in the lowest position contacting the substrate. This power stroke is largely amplified in the substrate-disengaging subunit (left panel), and is propagated counterclockwise through synchronization of three nucleotide-dependent events in three consecutive ATPases, i.e., ATP binding, ADP release and ATP hydrolysis (right panel).

Notably, the C-terminal tail of Rpt6 is inserted into the α2-α3 pockets in E_C1,2_ but not in E_B_ (Fig. 1g, Extended Data Fig. 4g). Curiously, the C-terminal tail of Rpt1, which is inserted in the α4-α5 pocket in E_B_, is instead unplugged from the α-pocket in E_C1,2_. Thus, the unplugging of Rpt1 C-tail from and the insertion of Rpt6 C-tail into the α-pockets appear to be synchronized with the disengagement of Rpt1 and engagement of Rpt6 with the substrate during the E_B_-to-E_C1,2_ transition, respectively.

Although the structure of AAA ring is nearly identical between states E_C1_ and E_C2_, multiple events are associated with the E_C1_-to-E_C2_ transition, including ADP release in Rpt1, ubiquitin dissociation from Rpn11, repositioning of the ATPase ring above the CP, and conformational changes in the lid (Fig. 4a, and Extended Data Fig. 4e,f and Table 2). Thus, states E_C1,2_ reflect the coordination between the initiation of substrate translocation and other regulatory events preparing the proteasome for processive substrate degradation.

## Power stroke

To understand the origin of the power stroke generated during nucleotide cycling that drives substrate translocation, we conducted a systematic structural comparison on the AAA domains of the ATPases. Structural alignment of the AAA domain of the same ATPase in different states against their large AAA subdomains defines a generic hinge-like rotation between the small and large AAA subdomains upon nucleotide binding (Figs. 3b,4b,5b). Rpt5 exhibits the greatest intrinsic hinge-like rotation of 25° between the small and large AAA subdomains during its engagement or disengagement with the substrate (Fig. 5a,b). By contrast, its three ADP-bound AAA structures in states E_A_ and E_C1,2_ are nearly identical to its two ATP-bound ones in E_B_ and E_D2_, suggesting the release of γ-phosphate after ATP hydrolysis is insufficient to drive intrinsic motion. Similarly, the dihedral angle between the small and large AAA subdomains of Rpt6 in its ADP-bound state in E_A_ and ATP-bound state in E_C1,2_ and E_D1,2_ remain nearly invariant, suggesting that nucleotide binding locks the AAA domain into a single rigid-body. Unlike Rpt5, Rpt2 assumes three categories of dihedral angles between its small and large AAA subdomains (Fig. 4b). The ATP-bound state in E_A_ and the ADP-bound state in E_B_ are similar, and are close to the apo-like state, but are quite different from the ATP-bound state in E_D1,2_. This likely reflects the observation that Rpt2 is the first one to disengage from the substrate after the initial substrate engagement with the AAA ring in E_B_. Nonetheless, in most cases, the major intrinsic conformational changes in the AAA domain of each ATPase occur between the apo-like and nucleotide-bound states. Thus, the most critical trigger for conformational change in the AAA domain of ATPases is either ATP binding or ADP release during substrate engagement or disengagement, respectively.

Next, to understand how the nucleotide-driven intrasubunit conformational change is amplified into a collective power stroke in the AAA ring, we align the entire AAA ring structures in different states against the large AAA subdomain of the ATPase that undergoes the complete cycle of nucleotide hydrolysis and exchange (Figs. 4c,d,5c-e). An out-of-plane rotation of 30° is found during the substrate-disengagement step of Rpt5 (Fig. 5c). Inspection of the AAA ring alignment in different viewing angle suggests that the local out-of-plane movement is more prominent in Rpt4 and Rpt3 than in Rpt1 and Rpt2, indicating that the intrinsic motion in Rpt5 is amplified to a greater degree along the counterclockwise direction than the opposite (Fig. 5c). By contrast, only 8° out-of-plane rotation is observed during the substrate-engagement step (Fig. 5d). The similar kind of unidirectional motion amplification is observed when the AAA subdomain of Rpt1, Rpt2, Rpt4 or Rpt6 is used for the AAA ring structure comparison (Fig. 4c,d). Thus, the collective power stroke in the AAA ring is largely amplified in the ATPase that is disengaging from the substrate from the bottom of the pore-loop staircase.

The above analysis allows us to postulate on a power stroke model integrating our structural observations. To generate processive power strokes, at least three consecutive ATPases must synchronize their nucleotide processing altogether: the first binding an ATP, the second releasing an ADP and the third hydrolyzing an ATP, as illustrated in Fig. 5g. Both ATP binding and ADP release trigger the similar degree of intrinsic hinge-like motion between their small and large AAA subdomains, but in opposite directions. Substrate disengagement in the second ATPase provides its AAA domain the degree of freedom to amplify the intrinsic hinge-like motions of both the first and second ATPases into a collective power stroke, resulting in the largest out-of-plane rotation (30-40°) in the substrate-disengaging ATPase. The power stroke facilitates ATP hydrolysis in the counterclockwise neighboring ATPase, likely through repositioning of the arginine fingers coordinating the neighboring ATP, which is essential for propagating the power stroke counterclockwise^41^. This power stroke makes room in the bottom that allows the other four substrate-bound ATPases to passively move down by differential out-of-plane rigid-body rotations of small angles (5-10°), which translate the substrate toward the CP through the pore-loop staircase.

## Long-range quaternary allosteric regulation

The proteasomal ATPase heterohexameric motor is asymmetrically regulated by Rpn1 and the lid subunits (Rpn5-7, 9). In both states E_D1_ and E_D2_, the Rpn1 toroidal domain forms a surface cavity with the CC domain of Rpt1-Rpt2, where a short helix from Rpn2 is inserted (Extended Data Fig. 7a-c)^42^. The short helix resides in the middle of a long loop (residue 820-871) emanating from Rpn2 toroidal domain. This long-range association of Rpn1-Rpn2 seems stabilize a larger interface formed between Rpn1-Rpn2 and Rpt1-Rpt2. However, such a quaternary architecture is not observed in other states (E_A-C_). In states E_C1,2_, the Rpn1 density is considerably blurred, reflecting strong motions potentially breaking the long-range Rpn1-Rpn2 associations seen in states E_D1,2_ (Fig. 1b, Extended Data Fig. 3a). This coincides with the nucleotide release in Rpt1-Rpt2 that facilitates the movement of Rpt6 to a higher position in the substrate-pore loop staircase. Thus, the specific Rpn1 conformation in each state appears to be highly coordinated with the ATP hydrolytic cycle in the AAA-ATPase and is controlled by its interactions with Rpn2 in a long-range fashion.

The Rpt4/5 CC domains also seem to act at a distance to help coordinate substrate translocation in a long-range fashion. In states E_C1,2_ and E_D1,2_, the N-terminal segment of Rpt4/5 CC domain interacts with the either Rpn9 or Rpn10 in different contacts (Extended Data Fig. 7d,e). By contrast, it does not contact any Rpn subunit in state E_A_ and E_B_, but instead, helps to recruit ubiquitin. Taken together, our data suggest that the lid regulates substrate interactions with the AAA-ATPase through a combination of short-range and long-range allosteric regulations.

## Insights into the complete cycle of substrate processing by 26S proteasome

Our data suggest that the ATP hydrolysis in the proteasome holoenzyme follows three distinct modes at different stages of substrate processing (Fig. 6). The first mode features ATP hydrolysis in a pair of oppositely positioned ATPases at a time. We speculate that this is orchestrated to promote initial substrate recognition and deubiquitylation, as evident in states E_A_ and E_B_ (Fig. 3d). This is reminiscent of the nucleotide-binding pattern observed in state S_D2_ of the substrate-free human proteasome and the caseinolytic protease X (ClpX) hexamer, an AAA-ATPase unfoldase in *E. coli*^26,43^. The second mode features two events of ATP hydrolysis in two consecutive ATPases at a time without advancing the substrate and is used to initiate the substrate translocation and to simultaneously regulate the CP gate opening (Fig. 4e). The third mode is greatly simplified as opposed to the other two modes and culminates with sequential hydrolysis of one nucleotide at a time (Fig. 5f). This is likely most efficient for maintaining processive substrate translocation, when the ubiquitin chain is removed, and the CP gate is open. In both the second and third modes, the synchronized ATP binding and ADP release in at least two consecutive ATPases can stimulate the power stroke of the AAA-ATPase ring that is largely amplified in the substrate-disengaging subunit and is propagated through the coordinated ATP hydrolysis in the counterclockwise adjacent ATPase. The third mode is reminiscent of the proposed ATP hydrolysis mechanism in several other hexameric ATPase motors^44-47^.

**Figure 6.**
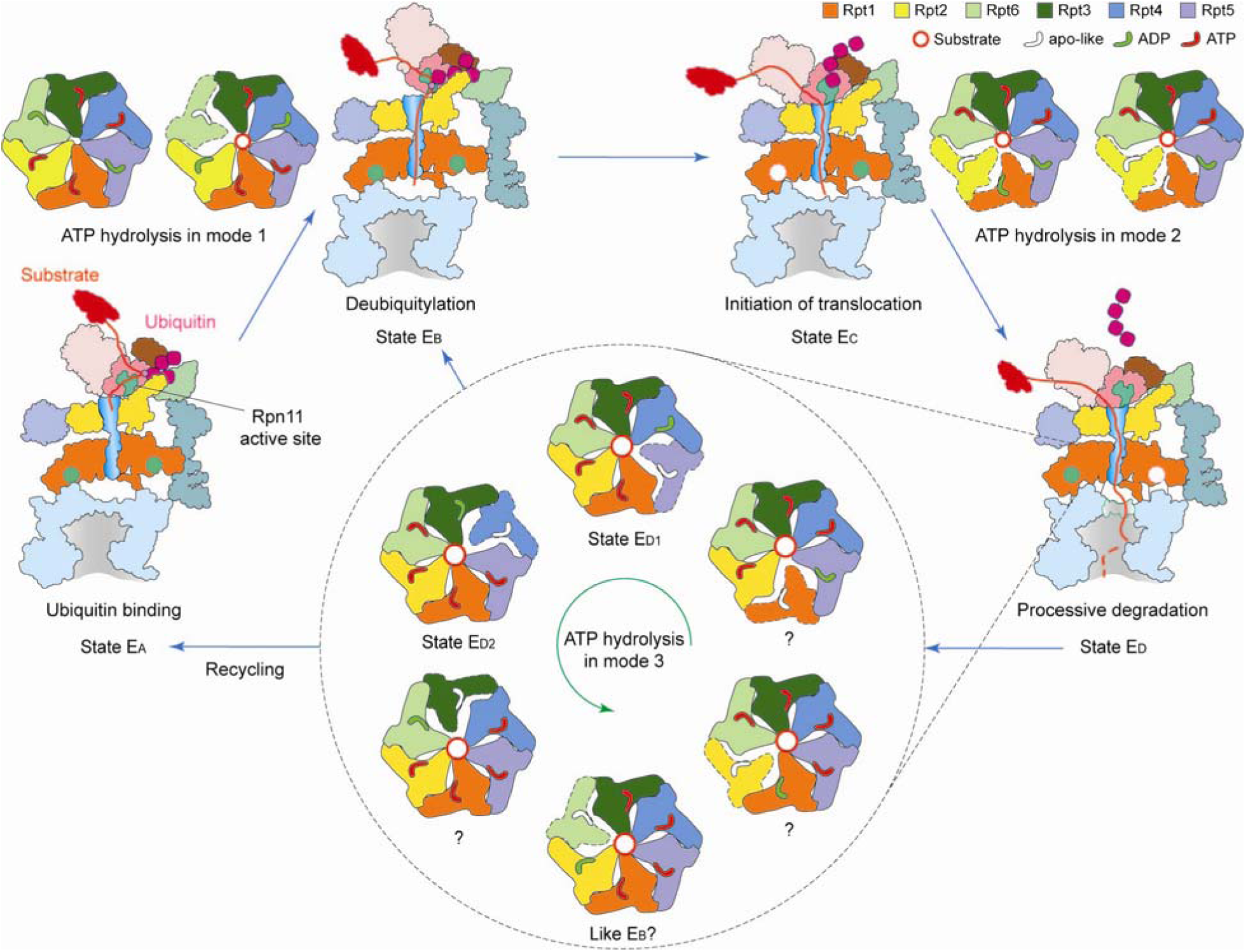
Insights into the complete cycle of substrate processing by the human 26S proteasome. The cartoon summarizes the concept of three principle modes of coordinated ATP hydrolysis observed in the seven states and explains how they regulate the complete cycle of substrate processing by the 26S holoenzyme. Coordinated ATP hydrolysis in modes 1, 2 and 3 features hydrolytic events in two oppositely positioned ATPases, in two consecutive ATPases and in only one ATPase at a time, respectively. The substrate processing undergoes three major steps before the CP gate opening for processive translocation: (1) ubiquitin recognition; (2) simultaneous deubiquitylation and substrate engagement with the AAA-ATPase ring; and (3) translocation initiation, which involves multiple simultaneous events, including ubiquitin release, ATPase repositioning and switching of Rpt C-tail insertion pattern. In steps 1 and 2, the ATPases follow the mode-1 ATP hydrolysis. In step 3, they follow the mode-2 ATP hydrolysis. After the gate is open, the AAA-ATPases follow the mode-3 ATP hydrolysis that only one nucleotide is hydrolyzed at a time.

Our data reveal the structural basis for the functional asymmetry of the six distinct ATPases^48-50^. Each ATPase is specialized for certain role in regulating substrate interactions that is complementary to the roles of the other ATPases. The broken symmetry accommodates the introduction of a distributed, quaternary allosteric regulation by the lid subcomplex, which sets “parity” on each ATPase essential for coordination between key steps, including substrate recognition, deubiquitylation, CP gating and processive translocation. Indeed, Rpt6 stands out to be uniquely essential to both the E_A_-to-E_B_ transition for substrate deubiquitylation and the E_C_-to-E_D_ transition for CP gate opening. The probability of assuming state E_D2_ is the highest among other translocation-compatible states. Rpt5 and Rpt1, whose AAA domain harbors no contacts with either the lid or Rpn1 are the ones at the highest position contacting substrate in state E_D2_. By contrast, the AAA domains of other four Rpt subunits all directly contact the lid subunits or Rpn1, which is also regulated by the lid through the long-range Rpn1-Rpn2 association.

In summary, we have determined the atomic structures of substrate-engaged human 26S proteasome in seven native states during its ATP-dependent degradation of polyubiquitylated substrate. These structures establish a foundation for understanding dynamic substrate-proteasome interactions during the complete cycle of substrate processing and provide a wealth of information accounting for several decades of biochemical studies of proteasome functions. A plethora of potential substrate-binding sites revealed in this study may facilitate future development of small-molecule inhibitors and drugs aiming at regulating the proteasomem functions implicated in various diseases, such as multiple myeloma and neurodegenerative diseases.

## Methods

### Preparation of polyubiquitylated Sic1^PY^

PY motif-inserted Sic1 (Sic1^PY^) and WWHECT were chosen as the model substrate and the E3 ubiquitin ligase, respectively. WWHECT was derived from wildtype Rsp5 with its N-terminal 220 amino acids deleted. Both of the proteins were expressed in *Escherichia coli* and purified as previously reported by Saeki et al.^30^, whose lab gifted the plasmids of Sic1^PY^ and WWHECT. After the transformation, BL21 (DE3) competent cells were grown to OD_600_ of 0.7 in LB medium with 50 μg/ml ampicillin when the cells were cooled to 30 °C and induced by 0.5 mM IPTG for 3 h. After being harvested (3000 g, 10 min), the cells were suspended and lysed by sonication in 50 mM PBS, pH 7.0, containing 300 mM NaCl, 10% glycerol, 1 mM DTT, 0.2% Triton X-100 and 1×protease inhibitor cocktail. The supernatant was recovered through the centrifugation (15,000 g, 30 min), which was then incubated with the pre-equilibrated TALON resin for 2 h at 4°C. After the binding, the resin was washed by 20 column of 50 mM Tris-HCl, pH 7.5, containing 100 mM NaCl, 10% glycerol, 1 mM DTT. The Sic1^PY^ was then eluted with the same buffer containing 150 mM imidazole. The elution was further purified by FPLC with buffer containing 50 mM Tris-HCl, pH 7.5, 100 mM NaCl, 10% glycerol through Superdex 75 at 0.25 mL/min.

To purify WWHECT, BL21 (DE3) competent cells were grown to OD_600_ of 0.5 in LB medium with 50 μg/ml ampicillin when the cells were cooled to 20 °C and induced by 0.2 mM IPTG for 15 h after the transformation. The cells were harvested and lysed through the same procedure as used for the Sic1^PY^. The supernatant was then incubated with the pre-equilibrated glutathione sepharose resin for 2 h at 4°C. After the binding, the resin was washed by 20 column of washing buffer (50 mM Tris-HCl, pH 7.5, containing 100 mM NaCl, 10% glycerol and 1 mM DTT) and then incubated with the same buffer containing PreScission protease for 12 h at 4°C. The resin was removed by centrifugation and the supernatant was then applied to FPLC through Superdex 75, with the same procedure descripted above.

To ubiquitinate the Sic1^PY^, 40 μg/ml Sic1^PY^, 500 nM Ube1 (Boston Biochem), 2 μM UbcH5a (Boston Biochem), 100 μg/ml WWHECT and 1 mg/mL ubiquitin (Boston Biochem) was incubated in the reaction buffer (50 mM Tris-HCl, pH 7.5, 100 mM NaCl, and 10% glycerol, 2 mM ATP, 10 mM MgCl2, and 1 mM DTT) for 3h at room temperature. Pre-equilibrated TALON resin was then incubated with the reaction system for 1 h at 4°C. After the resin was washed by 20-column volume of the wash buffer (50 mM Tris-HCl, pH 7.5, 100 mM NaCl, 10% glycerol), the polyubiquitinated Sic1^PY^ (PUb-Sic1^PY^) was eluted with the same buffer containing 150 mM imidazole. The elution was applied to Amicon ultra centrifugal filter with 30K molecular cut-off for removal of imidazole. The ubiquitination reaction was examined by Western blotting with anti-T7 antibody.

### Purification of the human 26S proteasome

Human proteasomes were affinity-purified on a large scale from a stable HEK293 cell line harboring HTBH (hexahistidine, TEV cleavage site, biotin, and hexahistidine) tagged hRPN11 (a gift from L. Huang, Departments of Physiology and Biophysics and of Developmental and Cell Biology, University of California, Irvine, California 92697)^51^. The cells were Dounce-homogenized in a lysis buffer (50 mM PBS, pH 7.5, 10% glycerol, 5 mM MgCl_2_, 0.5% NP-40, 5 mM ATP and 1 mM DTT) containing protease inhibitors. Lysates were cleared by centrifugation (20,000 × g, 30 min), then incubated with NeutrAvidin agarose resin (Thermo Scientific) for 3 h at 4 °C. The beads were then washed with excess lysis buffer followed by the wash buffer (50mM Tris-HCl, pH 7.5, 1mM MgCl_2_ and 1mM ATP). 26S proteasomes were cleaved from the beads by TEV protease (Invitrogen). The resin was removed by centrifugation and the supernatant was then further purified by gel filtration on a Superose 6 10/300 GL column at a flow rate of 0.15 ml/minute in buffer (30mM Hepes pH 7.5, 60 mM NaCl, 1 mM MgCl_2_, 10% glycerol, 0.5 mM DTT, 0.6 mM ATP). Gel-filtration fractions were concentrated to about 2 mg/ml and the buffer was exchanged to 50 mM Tris-HCl, pH 7.5, 100 mM NaCl, 1 mM ATP and 10% glycerol.

### Biochemical verification of the substrate-bound human proteasome

To verify the preparation of the PUb-Sic1^PY^, we performed degradation tests on the PUb-Sic1^PY^ using our purified human 26S proteasome. 100 nM proteasome was incubated with 20 μg/mL PUb-Sic1^PY^ for 10 min at 37°C in a buffer containing 50 mM Tris-HCl, pH 7.5, 100 mM NaCl, 10% glycerol, 5 mM ATP. The reaction was stopped by adding SDS loading buffer and 100 mM DTT. Samples were collected with the time points at 2 min, 5 min, and 10 min. Then the degradation reaction was examined by Western blotting with the anti-T7 antibody.

To verify the formation of proteasome-substrate complex in our cryo-EM imaging experiments, we crosslinked the proteasome-substrate complex and examined them with native gel electrophoresis. However, the crosslinking was not used for the sample preparation for cryo-EM data collection in order to preserve the native states of substrate interactions with the proteasome. Before crosslinking, the proteasome and PUb-Sic1^PY^ samples were first exchanged to a buffer containing 50 mM PBS, pH 7.5, 100 mM NaCl, 10% glycerol, 1 mM ATP by Zeba(tm) Micro Spin Desalting Columns (7K, Thermo Fisher). Then 1 μL 2 mg/mL PUb-Sic1 and 1 μL 1 mg/mL proteasome was mixed with 17 μL of the same buffer for 30s. After 1mM ATP-γS was added, 1 µl 2.3% freshly prepared solution of glutaraldehyde was added and incubated for 15 minutes at 37°C. The cross-linked complex was then examined by the native gel electrophoresis.

### Cryo-EM imaging and data collection

The 10 μL 2 mg/mL proteasome was incubated with 9 μL 2 mg/mL PUb-Sic1^PY^ for 30 s (50 mM Tris-HCl, pH 7.5, 100 mM NaCl, 10% Glycerol and 1 mM ATP) at room temperature and then 1 μL 20 mM ATP-γS was immediately added into the solution. To remove the glycerol, the complex system was applied to Zeba(tm) Micro Spin Desalting Columns (7K, Thermo Fisher) to exchange the buffer to 50 mM Tris-HCl, pH 7.5 containing 100 mM NaCl, 1mM ATP and 1 mM ATP-γS. Right before cryo-EM sample preparation, 0.005% NP-40 was added. Cryo-EM grids were prepared with FEI Vitrobot Mark IV. C-flat grids (R1/1 and R1.2/1.3, 400 Mesh, Protochips, CA, USA) were glow-discharged before a 2.5-μl drop of 1.5 mg/ml substrate-engaged proteasome solution was applied to the grids in an environment-controlled chamber with 100% humidity and temperature fixed at 4 °C. After 2 second blotting, the grid is plunged into liquid ethane and then transferred to liquid nitrogen. The cryo-grids were initially screened in an FEI Tecnai Arctica microscope, equipped with an Autoloader and an acceleration voltage of 200 kV at a nominal magnification of 235,000 times. Good quality grids were transferred to an FEI Titan Krios G2 microscope equipped with the post-column Gatan BioQuantum energy filter connected to Gatan K2 Summit direct electron detector. Coma-free alignment was manually optimized and parallel illumination was verified prior to data collection. Cryo-EM data were collected semi-automatically by the Leginon^56^ version 3.1 and SerialEM with the Gatan K2 Summit operating (Gatan Inc., CA, USA) in a super-resolution counting mode and with the Gatan BioQuantum operating in the zero-loss imaging mode. A total exposure time of 10 second with 250 ms per frame resulted in a 40-frame movie per exposure with an accumulated dose of 44 electrons/Å^2^. The calibrated physical pixel size and the super-resolution pixel size are 1.37 Å and 0.685 Å, respectively. The defocus in data collection was set in the range of −0.7 to −3.0 μm. A total of 44,664 movies were collected throughout eight long sessions of data collection.

### Cryo-EM data processing and reconstruction

The micrograph frames of 44664 raw movies were aligned and averaged with MotionCor2 program^52^. Each drift-corrected micrograph was used for the determination of the micrograph CTF parameters with program Gctf^17^. 2,669,687 particles of 26S proteasome were picked using program deepEM^53^. Reference-free 2D classification and 3D classification were carried out in both RELION 2.1^54^ and ROME, which combined the maximum-likelihood based image alignment and statistical machine-learning based classification^55^. Focused 3D classification, which we used in the later stage of data processing, and high-resolution refinement, were mainly done with RELION 2.1. Map reconstruction and local resolution calculation were finished with programs in both RELION 2.1 and ROME.

We applied a hierarchical 3D classification strategy to analyze such a huge dataset (Extended Data Fig. 2). The entire data-processing procedures include four steps. In the first step, we separated doubly capped proteasome particles from those singly capped ones, so as to generate pseudo-single-capped particles. All particles underwent several rounds of 2D and 3D classification batch by batch, resulting in 1,552,828 doubly capped particles and 478,919 singly capped ones. These particles were aligned to the consensus models of doubly and singly capped proteasomes to get their approximate shift and angular parameters. With these parameters, each complete double-capped particle was split into two pseudo-single-cap particles by re-centering the box onto the RP-CP subcomplex. Then the box size of pseudo-single-capped particles and true single-capped particles was shrunk to 600×600. This is an effective way to reduce the irrelevant heterogeneity due to conformational variations, and improve the map resolution^24,26^. There were totally 3,584,040 particles remained in the dataset for the following steps. In the second step, we focused on the gate of the CP. Likewise, several rounds of 2D and 3D classification were done to distinguish the states of the CP gate, that is, separating the S_A_-like closed-gate states from those open-gate or namely S_D_-like states. It is obvious that the RP subcomplex of the S_D_-like states rotates by a large angle compared to the S_A_-like states^15,26^. The RP-CP subcomplex was masked during the 3D classification. There were 732,666 particles in the S_A_-like states and 2,521,686 particles in the S_D_-like states left after this step. In the third step, we used focused 3D classification method^54^ to further classify within these two different states. Since the CP is structurally stable, we did refinement with the CP masked, so that we can get the x-y shift and angular parameters of all particles when they were aligned against the reference of the CP. Using these parameters, we continued 3D classification with the RP masked and with alignment skipped. That means we only classify the images based on their structural changes in the RP relative to the CP. After the classification, we can clearly see the RP dramatically swings and rotates against the CP. By focused on the variation of the substrate interactions with the AAA-ATPase and Rpn10/11, we classified these particles into 5 major states, designated E_A_, E_B_, E_C_, E_D1_, E_D2_, respectively, accounting for 7.8%, 14.8%, 9.9%, 27.2%, 40.1%. In the final step, we used focused classification to further detect significant structural changes within each of these five states. After auto-refinement with RP masked, we continued skip-alignment classification with the lid, the AAA-ATPase, or certain combination of RP subunits masked. Application of differential masks depended on specific structural characteristics of different states. For example, Rpn1 in state E_B_ state was partially blurred without further 3D classification. So we masked Rpn1 together with ATPase in 3D classification, which resulted in improvement of its density quality in certain 3D classes. The whole RP complex is highly dynamical in state E_C_. Thus, we did further classification with the whole RP masked, resulting in two distinct states, named E_C1_ and E_C2_. E_C1_ showed a clear ubiquitin density, which is absent in E_C2_. Similarly, we also obtained an intermediate state from initial E_B_ dataset, named E_A2_, which showed different ubiquitin-binding mode from that in E_A_.

The final refinement of each state was done on the particle data in the counting mode with a pixel size of 1.37 Å. Based on the in-plane shift and Euler angle of each particle from the last iteration of refinement, we reconstructed the two half-maps of each state using raw single-particle images at the super-counting mode with a pixel size of 0.685 Å. The final reconstructions of the E_A1_, E_A2_, E_B_, E_C1_, E_C2_, E_D1_, and E_D2_ datasets gave overall resolutions of 3.0 Å, 3.2 Å, 3.3 Å, 3.5 Å, 3.6 Å, 3.3 Å and 3.2 Å, respectively, measured by the gold-standard FSC at 0.143-cutoff on two separately refined half maps. Because states E_A1_ and E_A2_ exhibit identical structures in their CP and AAA-ATPase components, we combined them together and refined the combined dataset by applying the CP/ATPase mask, which yielded reconstructions measured to 2.8 Å resolution by the gold-standard FSC. To enhance the local density quality for each state, we applied two types of local mask in the last several iterations of refinement, one focusing on the complete RP and the other focusing on the CP and ATPase components, which yielded two maps for each state that showed improved local resolution in the lid and in the CP, respectively. Prior to visualization, all density maps were sharpened by applying a negative B-factor. Local resolution variations were further estimated using ResMap on the two half maps refined independently^56^.

### Atomic model building and refinement

The higher-resolution cryo-EM maps allowed us to refine atomic models with improved quality and to extend sequence registry beyond the published structures of the substrate-free proteasomes through *de novo* modeling (Extended Data Table 1). Because we did not stall the substrates in a homogeneous location during their degradation, the substrate densities were modelled using polypeptide chains without assignment of amino acid sequence to appreciate the nonspecific nature of substrate translocation in the 26S proteasome, except for the lysine residue forming a visible isopeptide bond with the ubiquitin in state E_B_.

To build the initial atomic model of the substrate-bound 26S proteasome complex, we used previously published human 26S structures as starting models and rebuilt each atomic model in Coot ^67^ for the seven conformational states. In states E_D1,2_, a number of residues at the N-terminal of Rpt1 and Rpt2 coiled coil domain were missing but was shown as reliable densities with flanking of large-side chains. These high-resolution features allowed us to conduct de novo tracing of these previously missing elements. In all previously published cryo-EM structures of human 26S proteasomes, the local resolution of the lid subcomplex was generally worse than 4.9 Å and was insufficient to ensure the correct registry of the side chains. Our density maps of all states, particularly E_B_ and E_D1,2_, exhibit significantly improved local resolution in the lid subcomplex (Extended Data Table 1 and Extended Data Fig. 3), allowing us to rebuild the majority of the lid subcomplex. The local resolution of Rpn1 in states E_B_ and E_D1,2_ also reached 4-5 Å, allowing us to improve the backbone model and make partial side-chain registry. Rpn1 have very poorer local resolutions in states E_C1,2_ and excluded atomic modelling at all. Thus, we used the improved atomic model of Rpn1 from states E_B_ and E_D1,2_ to fit the poor Rpn1 densities of E_C1,2_ only as a rigid-body.

The nucleotide densities are overall good for differentiating ADP from ATP, which allowed us to build the atomic models of ADP and ATP into their densities. A resolution of no worse than 3.6 Å may be required to distinguish ADP from ATP in nucleotide assignment into the density, because ATP adds an extra size of 2.46 Å with its γ-phosphate and three additional oxygens atoms relative to ADP. The magnesium ion bound to ATP was well resolved in state E_A_ (Extended Data Fig. 3c). By contrast, no magnesium ion density was observed around the ADP-assigned nucleotide density. Thus, the magnesium density was used to verify our ATP and ADP assignment. Except state E_A1,2_, at least one of the ATPases per state has a very poor nucleotide density quality in its nucleotide-binding site. Although there are visible extra densities in the nucleotide-binding site at a low contour level in most of the apo-like ATPase subunits after the protein structures are in place, these weak extra densities are insufficient for even fitting a complete ADP with good confidence. For instance, some of them may allow fitting of ribose and/or α-phosphate but then β-phosphate is totally out of density. To avoid over-interpretation and to practice prudence in high-quality atomic modeling, we avoided building atomic models of nucleotides into these poor densities at all and referred to the corresponding ATPases as the “apo-like state” throughout this study. The poor extra densities in the nucleotide-binding sites of these apo-like ATPases most likely reflect partial or low occupancy or unstable binding of nucleotide, which is expected when the nucleotide-binding site undergoes nucleotide exchange.

Atomic model refinement was conducted in Phenix with its real-space refinement program. We used both simulated annealing and global minimization option with NCS, rotamer and Ramachandran constraints. Partial rebuilding, model correction and density-fitting improvement in Coot were iterated after each round of atomic model refinement in Phenix^57^. The improved atomic models were then refined again in Phenix, followed by rebuilding in Coot^58^. The refinement and rebuilding cycle was repeated until the model quality reached the expectation (Extended Data Table 1).

### Structural analysis and visualization

All figures of the structures were plotted in Chimera^59^, PyMOL^14^, or Coot^58^. Structural alignment and comparison were performed in both PyMOL and Chimera. Interaction analysis between adjacent subunits was performed using PISA^60^.

### Data availability

The final cryo-EM maps and corresponding atomic models are in the process of being deposited into EM Data Bank and Protein Data Bank. They will be provided before formal publication of this manuscript.

## Acknowledgments

This work was funded in part by a grant of the Thousand Talents Plan of China, by an Intel academic grant, by the research funds at Peking-Tsinghua Center for Life Science at Peking University, by grants from National Natural Science Foundation of China 11774012, 91530321 (to Y.M.), and by the NIH grant GM43601 (to D.F.). The cryo-EM data were collected from the Cryo-EM Platform at Peking University. Initial cryo-EM screening experiments were performed in part at the Center for Nanoscale Systems at Harvard University, a member of the National Nanotechnology Coordinated Infrastructure Network (NNCI), which is supported by the National Science Foundation under NSF award no. 1541959 and the NIH grant AI100645. The data processing was performed in part in the Sullivan cluster, which is funded in part by a gift from Mr. and Mrs. Daniel J. Sullivan, Jr. and in the Weiming No.1 and Life Science No. 1 High-Performance Computing Platform at Peking University.

## Author contributions

Y.D. purified proteasome, prepared and screen the cryo-specimen condition for optimal data collection and conducted biochemical analysis. Y.D., S.Z, Z.W.,W.L.W., X.L. and Y.Z. and S.S.M. contributed to cryo-EM screening and data collection. S.Z., Z.W.,W.L.W. and Y.M. processed the data and refined the maps. Y.D., S.Z. and Y.M. built and refined the atomic models. Y.M. designed and supervised this study, formulated the concepts and drafted the manuscript. Y.D., S.Z., Y.L. and D.F. contributed to the manuscript preparation and structural analysis.

---

**Extended Data Figure 1.**
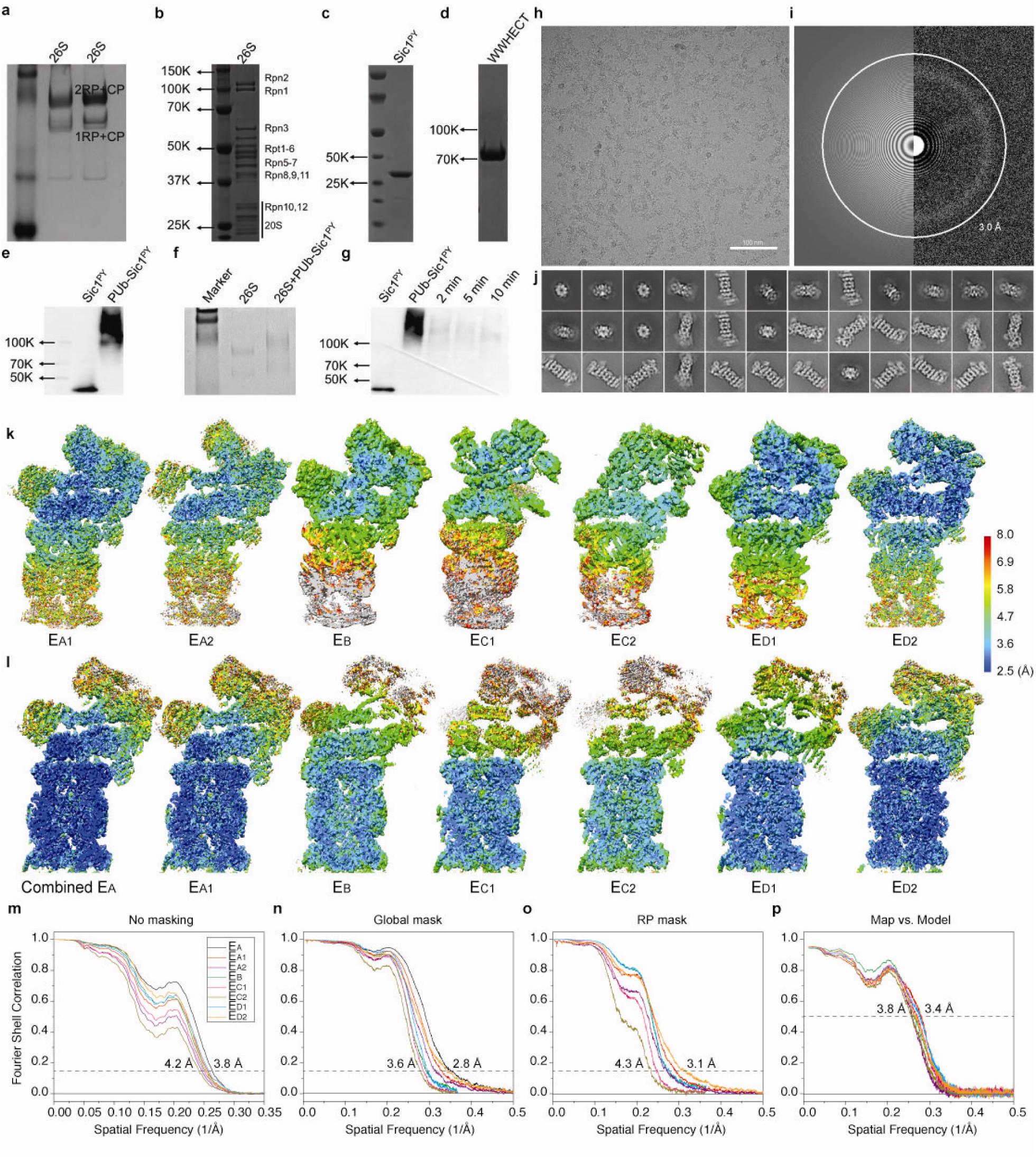
Characterization and structure determination of the substrate-engaged human proteasome. (**a**) Native PAGE analysis of proteasome purified through Superose 6 10/300 GL column. (**b**) SDS-PAGE results of the 26S proteasome after the FPLC. (**c)** SDS-PAGE analysis of the Sic1^PY^ purified through Superdex 75 column. (**d**) SDS-PAGE analysis of the WWHECT purified through Superdex 75 column. All PAGE from **a** to **d** were stained by coomassie blue. (**e**) Result of polyubiquitinated Sic1^PY^. After the ubiquitination, the samples were applied to SDS gel. Then the gel was examined by western-blot with anti-T7 antibody. The result suggests that almost all Sic1^PY^ has been ubiquitinated. (**f**) Native PAGE analysis of the crosslink between the proteasome and the substrate. As a control, 26S without substrate was also applied to the same crosslinking assay. With the addition of the PUb-Sic1^PY^, the proteasome was observed to run slower than those without PUb-Sic1^PY^. This result verifies that the substrate has been captured by the proteasome but not totally degraded yet at the time point when they were prepared for cryo-EM experiments. However, the crosslinking assay, designed only to verify the substrate-bound state of the proteasome, was not used in our cryo-EM sample preparation for data collection (see Methods). (**g**) SDS-PAGE analysis of the degradation of PUb-Sic1^PY^. The gel was examined by western-blot with anti-T7 antibody. Obvious degradation of the PUb-Sic1^PY^ has been observed, which suggests that both the proteasome and substrate were suitable for the degradation study. (**h**) A typical cryo-EM micrograph of the substrate-engaged human proteasome after motion correction. (**i**) The power spectrum evaluation of the micrograph shown in **h**. (**j**) A gallery of unsupervised class averages calculated by ROME^55^ using machine-learning-based clustering. (**k**) The local resolution estimation calculated by ResMap^56^ on seven maps refined by focusing the mask on the RP component. (**l**) The local resolution estimation on seven maps refined by focus the mask on the CP and ATPase components. (**m**) The gold-standard FSC plots of eight maps calculated without masking the raw half maps. (**n**) The gold-standard FSC plots of the maps refined with focusing on the CP and ATPase components, calculated with masking the raw half maps. (**o**) The gold-standard FSC plots of the maps refined with focusing on the RP subcomplex, calculated with masking the raw half maps. (**p**) The cross-map FSC plots calculated by Phenix between the each refined map and its corresponding atomic model. For each state, the maps refined by differential masking were merged in Fourier space into a single map, which was used for the cross-map FSC calculation. The same color code is used in panels **m**-**p**.

**Extended Data Figure 2.**
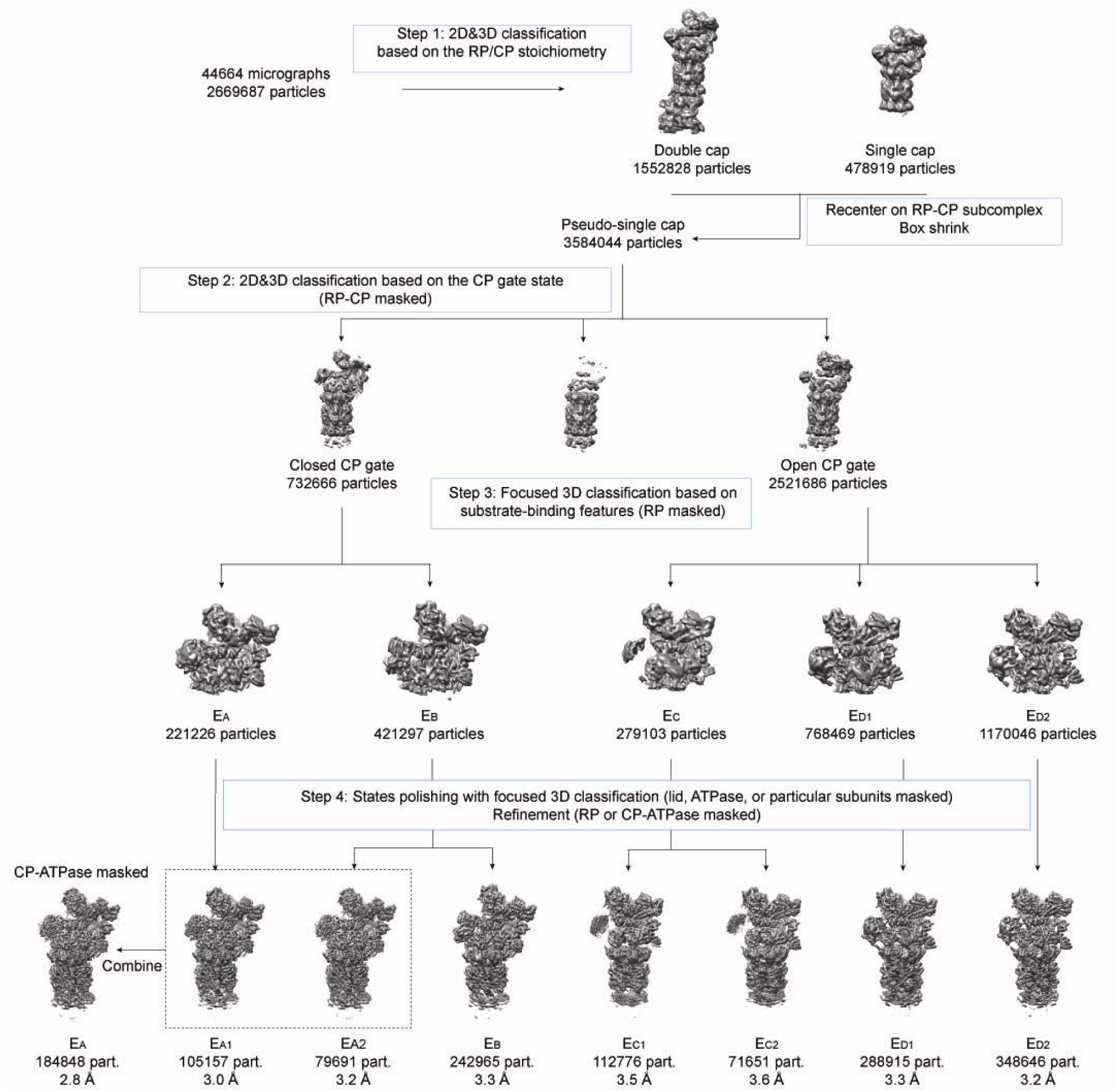
Focused classification to sort out the seven conformational states. The diagram illustrates the major four steps of our hierarchical focused classification strategy. Further detailed iterations of classification in each step are omitted for clarify.

**Extended Data Figure 3.**
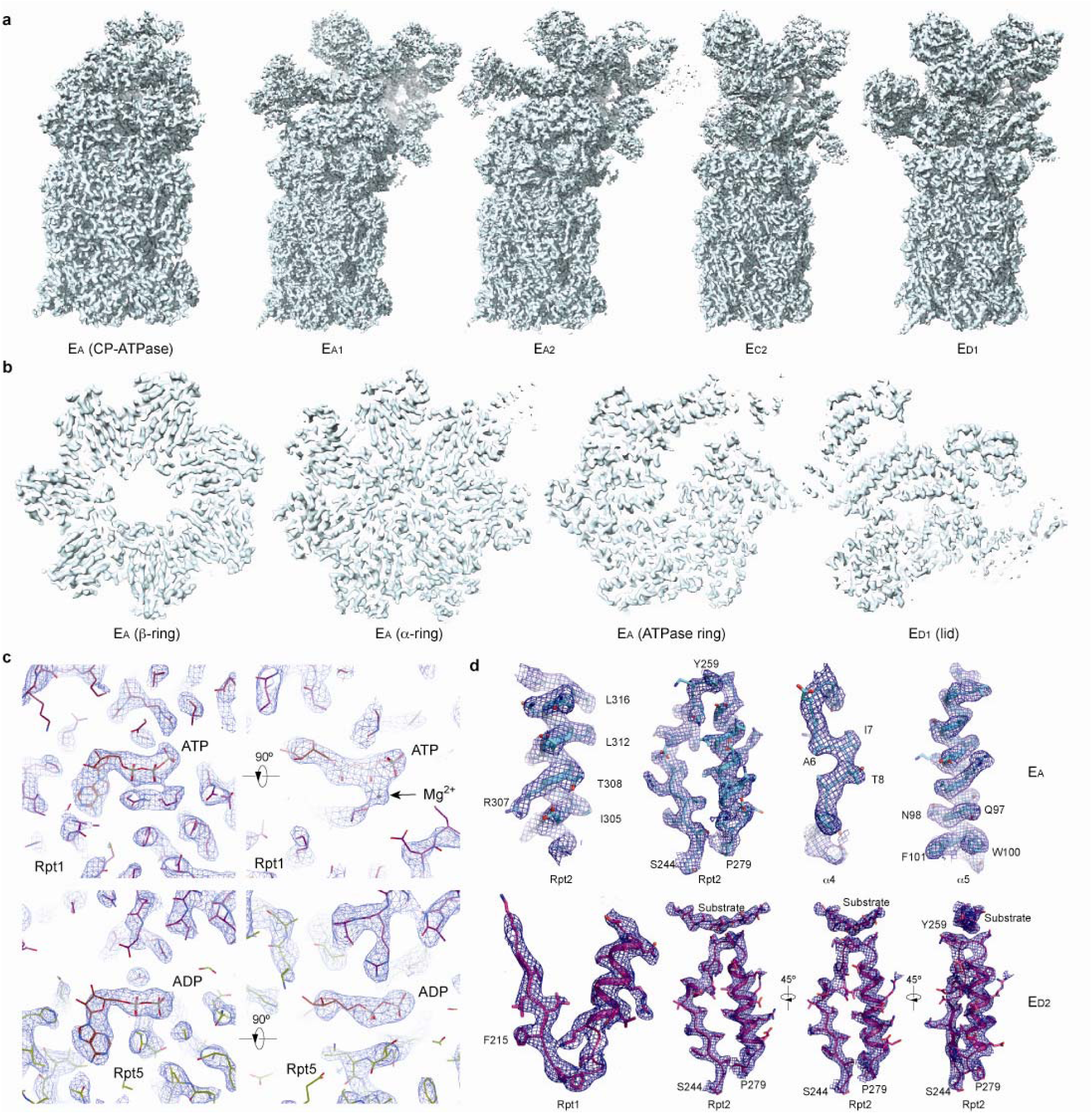
The cryo-EM maps and quality assessment. (**a**) The other five refined cryo-EM maps that are not shown in the main figures. (**b**) Typical central cross-sections in the density maps for each of the four subcomplexes, the lid in state E_D1_, the ATPase ring, the α ring and the β ring in state E_A_. (**c**) Typical nucleotide-binding pocket densities from the E_A_ map. The ATP density is compared with the ADP density in two orthogonal perspectives. The magnesium density next to the ATP is labeled, which is absent in the ADP-bound pocket. (**d**) Typical densities of secondary structures and substrate in the proteasome superimposed with their atomic models.

**Extended Data Figure 4.**
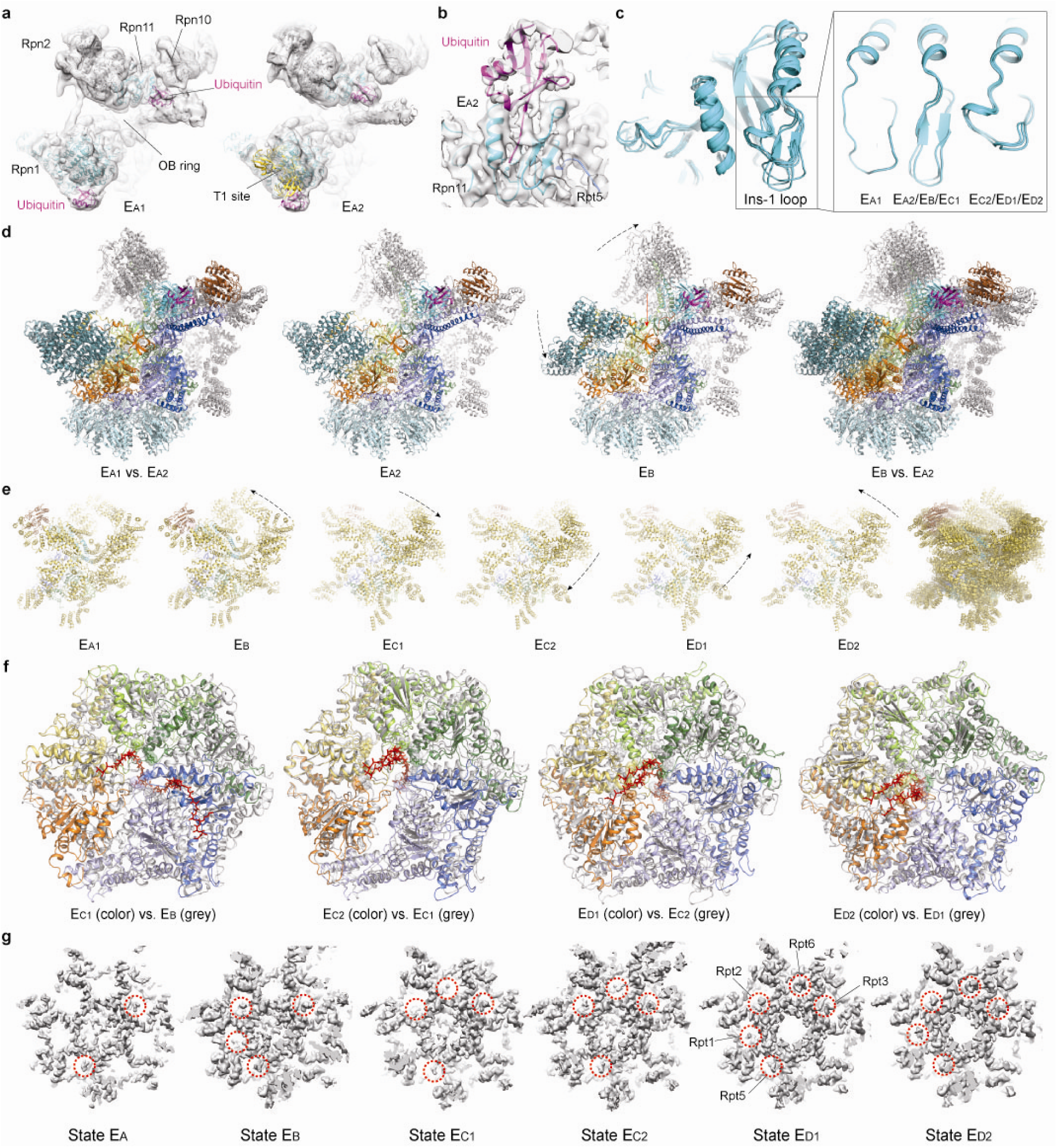
Key structural features and comparisons that help sort out the sequence of the seven conformational states. (**a**) The ubiquitin densities in state E_A1_ (left) and E_A2_ (right). The T1 site is labelled by fitting the yellow cartoon representation of the NMR structure (PDB ID 2n3u) of Rpn1 T1 element in complex with two ubiquitins into our density, showing the ubiquitin on Rpn1 is bound to a site very close to the T1 site^17^. The density maps are low-pass filtered to 8 Å to show the ubiquitin features clearly, due to the lower local resolution of the ubiquitin density in these maps. (**b**) The ubiquitin-Rpn11-Rpt5 interface observed at high resolution in state E_B_ is also observed in state E_A2_, albeit at somewhat lower local resolution. The E_A2_ density is shown as a transparent surface. (**c**) Comparison of the Insertion-1 loop of Rpn11 in different states. (**d**) Comparison of the RP structures between E_A_ and E_B_. (**e**) Comparison of the lid subcomplex conformations among all states. (**f**) Comparison of ATPase ring structures between two successive states. The structures are aligned together against their CP in **d**-**f**. (**g**) Comparison of the RP-CP interface and the Rpt C-tail insertions into the CP surface pockets in different states. The cryo-EM densities of the RP-CP interfaces are shown as a grey surface representation.

**Extended Data Figure 5.**
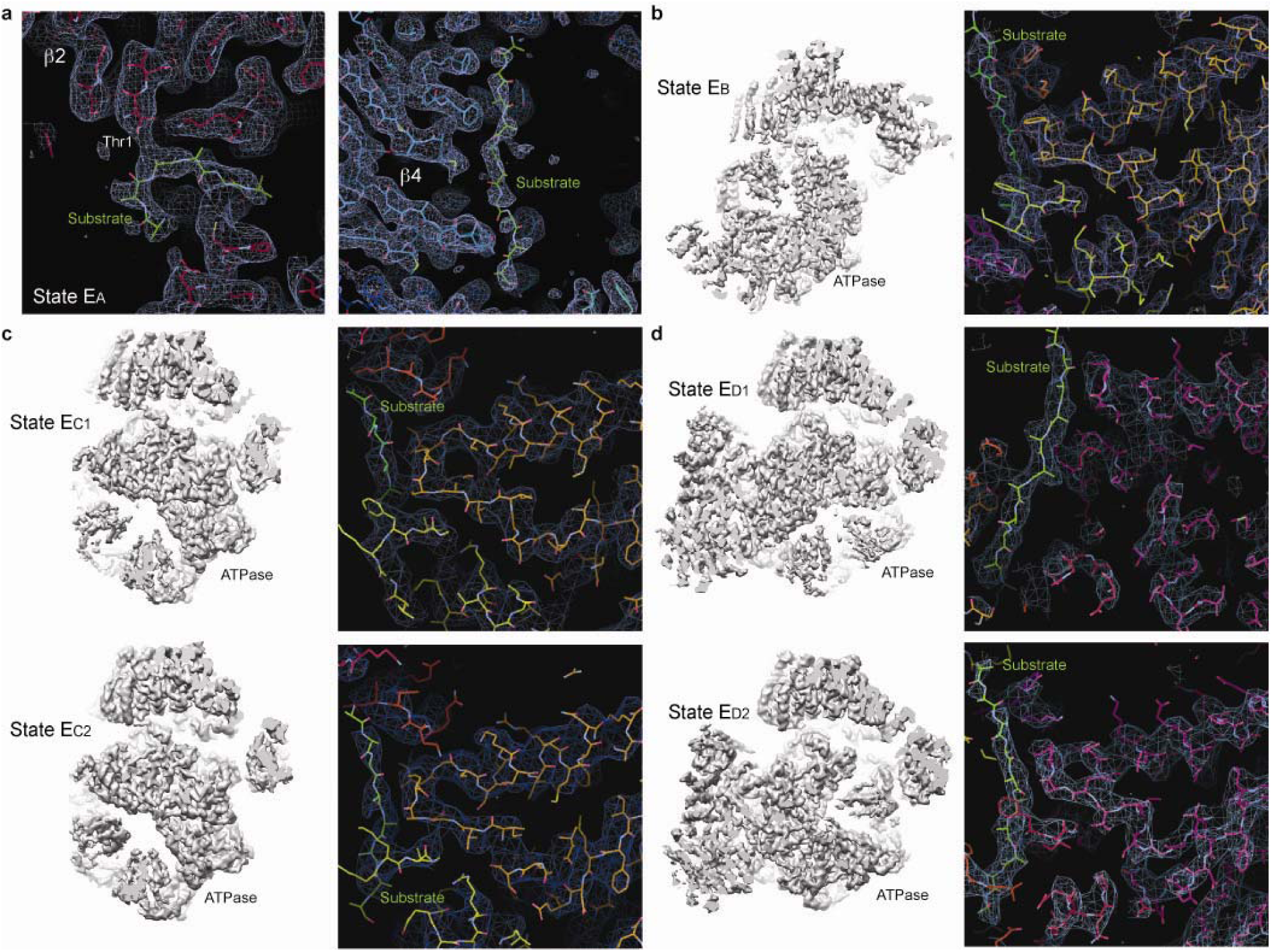
Substrate densities in different states. (**a**) Close-up views of two typical substrate densities observed in the CP chamber in state E_A_. The left panel shows the substrate density directly contacting the proteolytically active Thr1 in the β2 subunit. The right panel shows a long substrate density at the seam between two β4 subunits inside the CP. (**b**) The overall ATPase ring density of state E_B_ (left) and a close-up view of the substrate density (right). (**c**) The overall ATPase ring density of state E_C1,2_ (left) and a close-up view of the substrate density (right). (**d**) The overall ATPase ring density of state E_D1,2_ (left) and a close-up view of the substrate density (right). All close-up views were directly screen-copied from Coot^58^ after atomic modelling into the density maps without further modification or beautification.

**Extended Data Figure 6.**
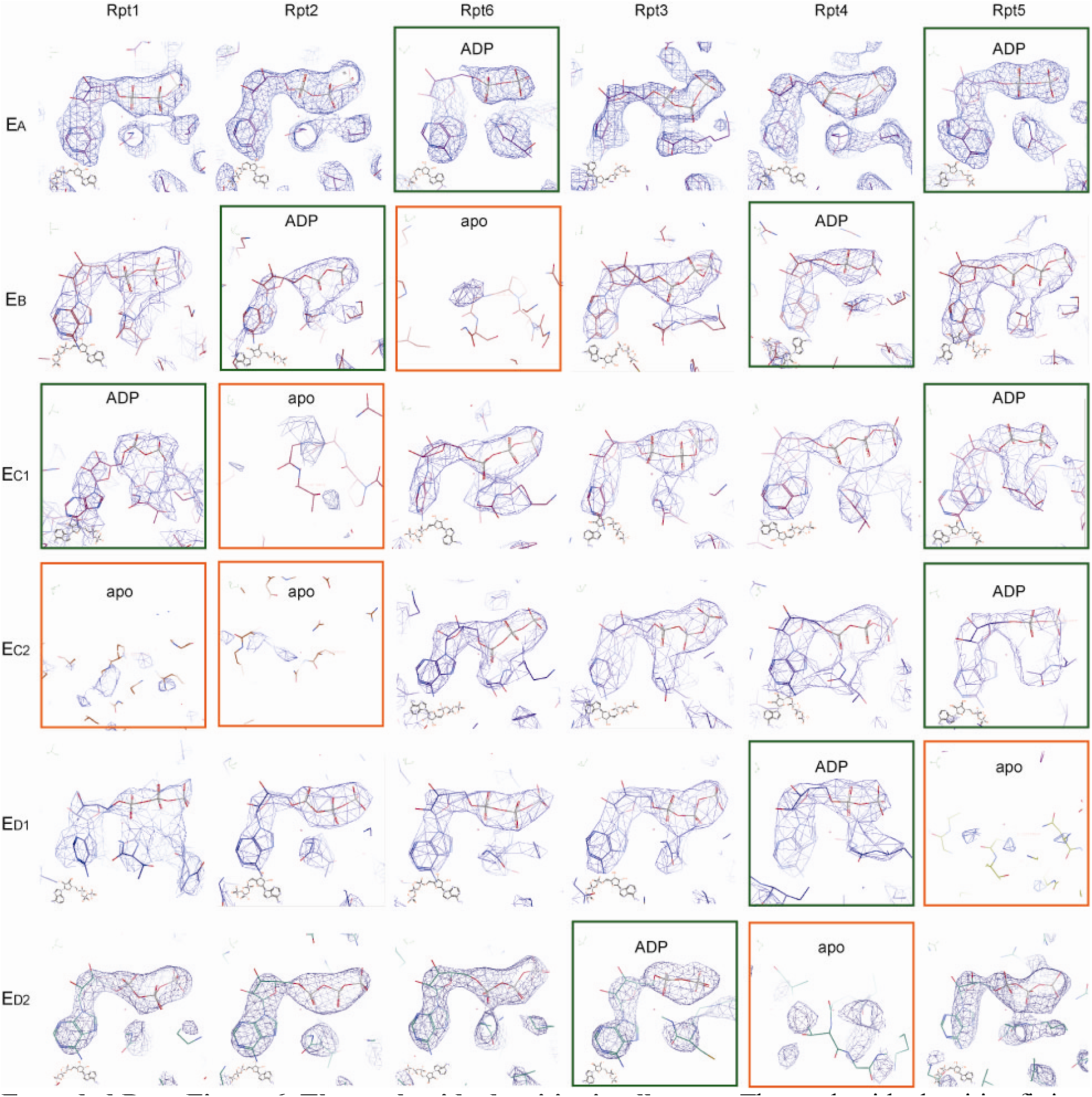
The nucleotide densities in all states. The nucleotide densities fitting with atomic models are shown with blue meshes. All close-up views were directly screen-copied from Coot^58^ after atomic modelling into the density maps without further modification or beautification. At the common contour level used for atomic modelling, the potential nucleotide densities in the apo-like subunits are mostly out, albeit they can show up partial nucleotide shapes at much lower contour level.

**Extended Data Figure 7.**
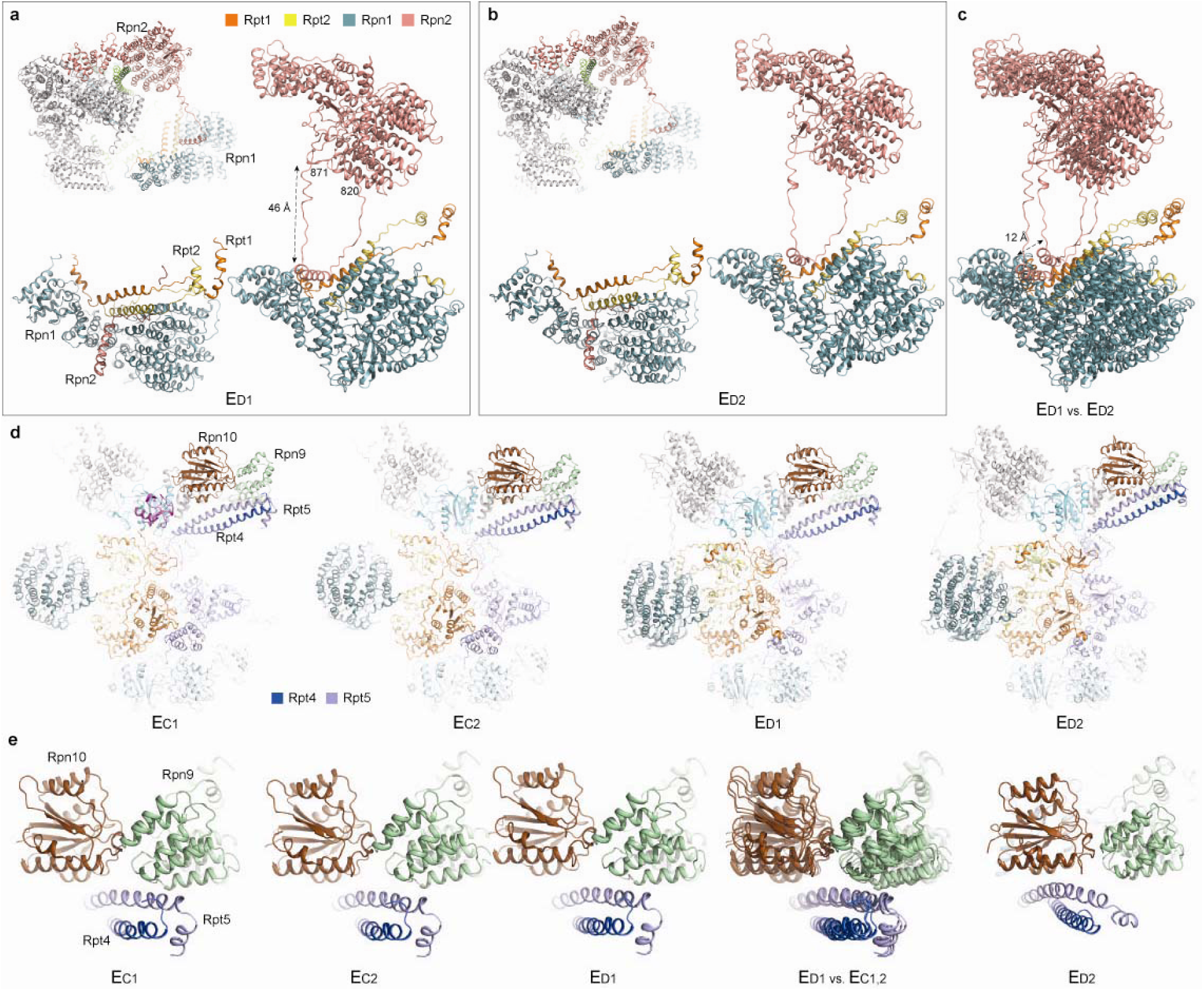
Long-range regulation of the AAA-ATPase by the lid-base interactions. (**a**) and (**b**) The long-range association of the Rpn1-Rpn2 through a looping structure from Rpn2 (residue 820-871) observed in state E_D1_ (**a**) and E_D2_ (**b**). (**c**) Comparison of the Rpn1-Rpn2 long-range association between the two states shows a marked movement of 12 Å. (**d**) The comparison of the Rpt4-Rpt5 CC interactions with Rpn9 and Rpn10 in states E_C1,2_ and E_D1,3_. (**e**) Closeup views of the Rpt4-Rpt5 CC contacts with Rpn9 in states E_C1,2_ and E_D1_, and its contact switching to Rpn10 in state E_D2_.

**Extended Data Table 1.** Data collection, structure determination and refinement statistics.

| Cryo-EM data collection statistics |  |  |  |  |  |  |  |  |
| --- | --- | --- | --- | --- | --- | --- | --- | --- |
| Microscope | Titan Krios G2 operating at 300 kV acceleration voltage with parallel beam |  |  |  |  |  |  |  |
| Camera | Gatan K2 Summit in the super-resolution counting video mode |  |  |  |  |  |  |  |
| Energy filter | Gatan BioQuantum energy filter in the zero-loss imaging mode |  |  |  |  |  |  |  |
| Energy slit | 20 eV |  |  |  |  |  |  |  |
| Magnification | 105,000 |  |  |  |  |  |  |  |
| Total dose | 44 electron/ Å<sup>2</sup> |  |  |  |  |  |  |  |
| Dose rate | 4.4 electron / Å<sup>2</sup> / second |  |  |  |  |  |  |  |
| Frame per movie | 40 |  |  |  |  |  |  |  |
| Physical pixel size | 1.37 Å |  |  |  |  |  |  |  |
| Super-resolution pixel size | 0.685 Å |  |  |  |  |  |  |  |
| Total micrographs | 44,664 |  |  |  |  |  |  |  |
| Total particles | 2,669,687 (double caps), 3,584,040 (re-boxed pseudo-single caps) |  |  |  |  |  |  |  |
| Structure determination statistics |  |  |  |  |  |  |  |  |
| State | E<sub>A</sub> | E<sub>A1</sub> | E<sub>A2</sub> | E<sub>B</sub> | E<sub>C1</sub> | E<sub>C2</sub> | E<sub>D1</sub> | E<sub>D2</sub> |
| Final particles | 184,848 | 105,157 | 79,691 | 242,965 | 112,776 | 71,651 | 288,915 | 348,646 |
| Overall resolution (Å) | 2.8 | 3.0 | 3.2 | 3.3 | 3.5 | 3.6 | 3.3 | 3.2 |
| Average resolution in the CP (Å) | 2.6 | 2.8 | 3.0 | 3.2 | 3.3 | 3.4 | 3.1 | 2.9 |
| Average resolution in the AAA-ATPase (Å) | 3.0 | 3.2 | 3.4 | 3.4 | 3.6 | 3.8 | 3.4 | 3.3 |
| Average resolution in the lid (Å) | N/A | 3.5 | 3.8 | 3.6 | 4.0 | 4.3 | 3.6 | 3.5 |
| Rotational accuracy (°) | 0.46 | 0.442 | 0.454 | 0.707 | 0.675 | 0.571 | 0.594 | 0.571 |
| Translational accuracy | 0.315 | 0.312 | 0.324 | 0.499 | 0.4 | 0.4 | 0.399 | 0.4 |
| Symmetry | C1 | C1 | C1 | C1 | C1 | C1 | C1 | C1 |
| B-factor (Å<sup>2</sup>) | -50 | -50 | -46 | -35 | -30 | -50 | -35 | -30 |
| Atomic model refinement statistics |  |  |  |  |  |  |  |  |
| Model-to-map CC (mask/volume) | 0.829/0.824 | 0.834/0.828 | 0.827/0.822 | 0.815/0.812 | 0.736/0.734 | 0.788/0.784 | 0.852/0.846 | 0.861/0.855 |
| RMSD Bond Length (Å) | 0.007 | 0.01 | 0.012 | 0.006 | 0.005 | 0.004 | 0.007 | 0.007 |
| RMSD Bond angles | 0.925 | 1.075 | 1.013 | 1.025 | 0.871 | 0.901 | 1.266 | 1.187 |
| Ramachandran favored (%) | 93.07 | 92.45 | 91.59 | 90.83 | 92.56 | 91.46 | 90.13 | 89.61 |
| Ramachandran outliers (%) | 0.17 | 0.22 | 0.26 | 0.34 | 0.23 | 0.23 | 0.29 | 0.3 |
| Rotamer outliers (%) | 0.34 | 0.42 | 0.42 | 0.66 | 0.28 | 0.19 | 0.45 | 0.5 |
| Clashscore | 2.69 | 2.91 | 5.02 | 5.63 | 4.00 | 4.2 | 5.87 | 5.83 |
| Molprobit score | 1.5 | 1.69 | 1.77 | 1.83 | 1.65 | 1.71 | 1.86 | 1.88 |

**Extended Data Table 2.** Summary of key structural features.

| **Key structural features connecting the seven states into a continuum of dynamic structural changes** |  |  |  |  |  |  |  |
| --- | --- | --- | --- | --- | --- | --- | --- |
| **States** | **E<sub>A1</sub>** | **E<sub>A2</sub>** | **E<sub>B</sub>** | **E<sub>C1</sub>** | **E<sub>C2</sub>** | **E<sub>D1</sub>** | **E<sub>D2</sub>** |
| Locations of ubiquitin densities | Rpn1, Rpn10, Rpt4/5 CC | Rpn11, Rpn10, Rpn1, Rpt4/5 CC | Rpn11, Rpn10, Rpt4/5 CC | Rpn11 | None observed | None observed | None observed |
| Rpn11 Ins-1 conformation | Large loop | β-hairpin | β-hairpin | β-hairpin | Retracted small loop | Retracted small loop | Retracted small loop |
| Pore-loop staircase contacting substrate | None | None | Rpt3 at the top, followed by Rpt4/5/1/2 | Rpt6 at the top, followed by Rpt3/5/1 | Rpt6 at the top, followed by Rpt3/5/1 | Rpt1 at the top, followed by Rpt2/6/3/4 | Rpt5 at the top, followed by Rpt1/2/6/3 |
| Rpt subunits disengaged from pore loop staircase | Rpt6 | Rpt6 | Rpt6 | Rpt1 and Rpt2 | Rpt1 and Rpt2 | Rpt5 | Rpt4 |
| Rpt subunits with apo-like state | None | None | Rpt6 | Rpt2 | Rpt1 and Rpt2 | Rpt5 | Rpt4 |
| Rpt subunits with ADP bound | Rpt6 and Rpt5 | Rpt6 and Rpt5 | Rpt2 and Rpt4 | Rpt1 and Rpt5 | Rpt5 | Rpt4 | Rpt3 |
| Rpt C-tails insertion into α-pockets | Rpt3 and Rpt5 | Rpt3 and Rpt5 | Rpt1/2/3/5 | Rpt2/3/5/6 | Rpt2/3/5/6 | Rpt1/2/3/5/6 | Rpt1/2/3/5/6 |
| CP gate state | Closed | Closed | Closed | Closed | Closed | Open | Open |
| Resemblance to the substrate-free conformations | S<sub>A</sub> | S<sub>A</sub> (in ATPase and CP) | None | S<sub>B</sub> (in overall lid-base relationship) | S<sub>C</sub> (in overall lid-base relationship) | S<sub>D</sub> (in overall lid-base relationship, RP-CP interface and CP gate) | S<sub>D2</sub> (in pore-loop staircase, overall lid-base relationship, RP-CP interface and CP gate) |
| **Potential substrate-binding sites observed in the CP chamber** |  |  |  |  |  |  |  |
| **State** | **Key contacts at the binding site** |  |  |  | **Features** |  | **Number of substrate residues** |
| E<sub>A</sub>, E<sub>B</sub>, E<sub>C1,2</sub>, E<sub>D1,2</sub> | Thr1, Cys31 of β2, Cys129 of β3 |  |  |  | Symmetric |  | 6 |
| E<sub>A</sub> | Asn24, Tyr134, Phe137 of β4 |  |  |  | At the seam between two β4 subunits |  | 10 |
| E<sub>A</sub> | Tyr103 of α1, Tyr61, Tyr90 of β1, and Phe88 of β2 |  |  |  | Symmetric, at the inter-subunit interface |  | 5 |
| E<sub>A</sub> | Asn90 of α5, Tyr90 of β5, Phe101 of β6 |  |  |  | Symmetric, at the inter-subunit interface |  | 3 |
| E<sub>A</sub>, E<sub>B</sub>, E<sub>C1,2</sub>, E<sub>D1,2</sub> | Tyr105, Arg117 of α1, His88 of α2 |  |  |  | Symmetric |  | 3 |
| E<sub>A</sub>, E<sub>B</sub>, E<sub>C1,2</sub>, E<sub>D1,2</sub> | Tyr59, Cys91, Tyr98 of β4 |  |  |  | Symmetric |  | 3 |
| E<sub>A</sub> | Phe69, Cys91 and Tyr98 of β3 |  |  |  | Asymmetric, only present in the chamber with the CP gate open |  | 3 |
| E<sub>A</sub> | Ile3, Tyr6, Tyr104 of β3, Tyr120 of β4 |  |  |  | Asymmetric, only present in the chamber with the CP gate open |  | 3 |
| E<sub>B</sub>, E<sub>C1,2</sub>, E<sub>D1,2</sub> | Tyr30 of β7 |  |  |  | Symmetric |  | 4 |

